# An Ancestral Function of Strigolactones as Symbiotic Rhizosphere Signals

**DOI:** 10.1101/2021.08.20.457034

**Authors:** Kyoichi Kodama, Mélanie K. Rich, Akiyoshi Yoda, Shota Shimazaki, Xiaonan Xie, Kohki Akiyama, Yohei Mizuno, Aino Komatsu, Yi Luo, Hidemasa Suzuki, Hiromu Kameoka, Cyril Libourel, Jean Keller, Keiko Sakakibara, Tomoaki Nishiyama, Tomomi Nakagawa, Kiyoshi Mashiguchi, Kenichi Uchida, Kaori Yoneyama, Yoshikazu Tanaka, Shinjiro Yamaguchi, Masaki Shimamura, Pierre-Marc Delaux, Takahito Nomura, Junko Kyozuka

## Abstract

In flowering plants, carotenoid-derived strigolactones (SLs) have dual functions as hormones that regulate growth and development, and as rhizosphere signaling molecules that induce symbiosis with arbuscular mycorrhizal (AM) fungi. Here, we report the identification of bryosymbiol (BSB), a previously unidentified SL from the bryophyte *Marchantia paleacea*. BSB is also found in vascular plants, indicating that it is ancestral in land plants. BSB synthesis is enhanced at AM symbiosis permissive conditions and BSB deficient mutants are impaired in AM symbiosis. In contrast, the absence of BSB synthesis has little effect on the growth and gene expression. We show that the introduction of the SL receptor of Arabidopsis renders *M. paleacea* cells BSB-responsive. These results suggest that BSB is not perceived by *M. paleacea* cells due to the lack of cognate SL receptors. We propose that SLs originated as AM symbiosis-inducing rhizosphere signaling molecules and were later recruited as plant hormone.

## INTRODUCTION

Land plants evolved from an algal ancestor more than 450 million years ago^1^. The first land plants faced many challenges, such as UV radiation and nutrient-poor soil. Thus, initial colonization of the terrestrial environment required the evolution of innovations such as the deployment of complex hormonal regulations and the mutualistic symbiosis formed with arbuscular mycorrhizal (AM) fungi, a monophyletic group of symbiotic soil-borne fungi^2, 3^. During AM symbiosis, plants produce and supply carbohydrates and lipids to the fungus, in exchange for mineral nutrients, in particular phosphorus, mined in the soil by the fungal symbiont^4^. Symbiosis with AM fungi is observed in more than 80% of extant land plant species, including the bryophytes that diverged from the vascular plant lineage more than 400 million years ago^3^. In flowering plants, establishment of the AM symbiosis requires the activation of fungal metabolism and stimulation of hyphal branching by plant root-derived strigolactones (SLs)^5^. In addition to their role as rhizosphere signaling molecules, SLs also function as a class of plant hormones and regulate various aspects of growth and development in flowering plants^6, 7^. The dual function, as hormones and as rhizosphere signaling molecules, makes SLs unique among plant hormones.

The function and signaling of SLs are well understood in flowering plants, whereas little is known outside this plant lineage^8, 9^. The origin and evolution of their dual function are also largely unknown. One reason why SL research has been hampered is related to their nature. The chemical structure of SLs differs from species to species, many species have multiple SLs and they exist in cells in minute concentrations^10, 11^. Phylogenetic studies and genome sequences revealed that genes in the SL biosynthesis pathway are conserved in bryophytes^12, 13^. However, although initial studies reported SLs in some bryophytes, their actual presence is still under debate and, more importantly, their function remains unknown^11, 12, 14, 15^.

Here we report the identification of an ancestral SL, bryosymbiol (BSB), present in diverse bryophytes such as *Marchantia paleacea*, and vascular plants. In *M. paleacea*, BSB is secreted from the plants and is required for AM symbiosis, but not for development. We show that BSB is not perceived by *M. paleacea* cells due to the absence of cognate SL receptors. Our findings reveal that the ancestral function of SLs is as AM symbiosis-inducing rhizosphere signaling molecules and that this function was already present in the most recent common ancestor of land plants.

## RESULTS

### Presence of SL biosynthesis genes is specific to AM symbiosis-forming species in *Marchantia*

Among the three extant groups of bryophytes, AM symbiosis is observed in liverworts and hornworts, but not in mosses^3, 16^. In liverworts, hyphae of AM fungi enter through rhizoids and produce arbuscules in parenchymatous cells in the midrib region of the thallus (Fig. 1a). We confirmed that AM symbiosis is widely observed in the genus *Marchantia*, including *M. paleacea, M. emarginata* and *M. pinnata*, whereas it is absent in *M. polymorpha*, the model liverwort species used for molecular genetic studies (Fig. 1a).

**Figure 1.**
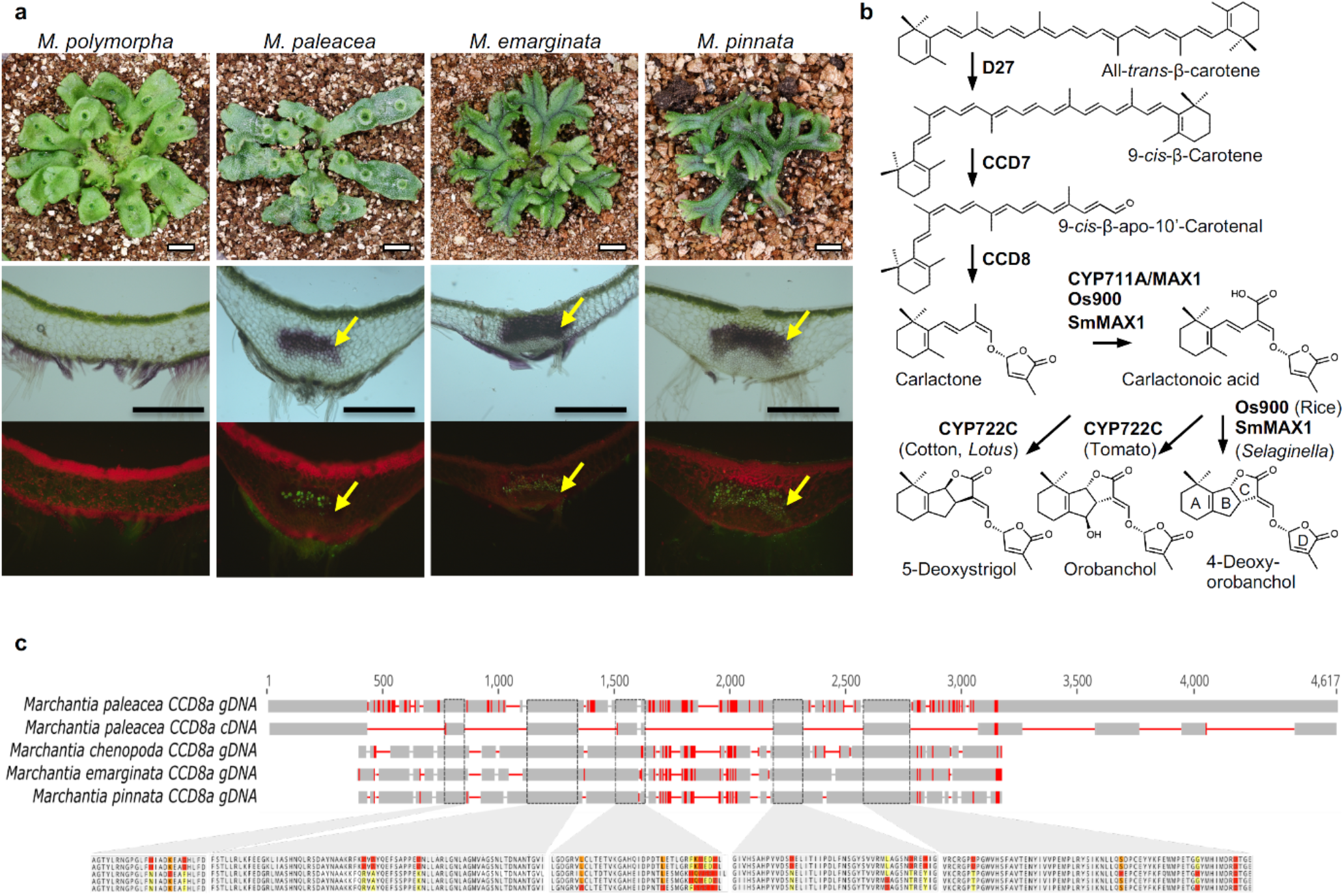
Presence of SL biosynthesis genes is specific to AM symbiosis-forming species in *Marchantia*. **(a)** *Marchantia* species, namely *Marchantia polymorpha, Marchantia paleacea, Marchantia emarginata* and *Marchantia pinnata*, used in this study Upper panels show plant morphology Scale bars: 1 cm. Middle and lower panels are transverse sections after colonization by *Rhizophagus irregularis*. The lower panel shows WGA-FITC staining of the fungi. Arbuscules are formed in the midrib region in the thallus (arrows). Scale bars: 0.5 mm. **(b)** Biosynthesis pathway for strigolactones in seed plants and lycophytes. **(c)** Conservation of the CCD8A genes in mycorrhizal Marchantia species. Amplified sequences of CCD8A are aligned against the *Marchantia paleacea* gene model. Red lines indicate polymorphisms. The predicted amino acid sequences in the conserved domains are shown below the gene model. Polymorphic amino acids are colored.

To date, more than 30 SLs have been identified in root exudates of various plant species^11^. SL biosynthesis starts from the conversion of all-*trans*-β-carotene to 9-*cis*-β-carotene by DWARF27 (D27)^17^ (Fig. 1b). Then, 9-*cis*-β-carotene is converted to carlactone (CL) by the consecutive actions of two carotenoid cleavage dioxygenases (CCD7 and CCD8)^18, 19, 20^. Carlactone is converted to carlactonoic acid (CLA), the universal precursor of a variety of species-dependent SLs, by the cytochrome P450, CYP711A, encoded by *MORE AXILLARY GROWTH 1* (*MAX1*) homologs^21^. Carlactonoic acid is converted to canonical SLs such as 4-deoxyorobanchol (4DO), orobanchol and 5-deoxystrigol (5DS) by species-specific functions of the P450s, CYP711A and CYP722C^22, 23^. The *M. paleacea* genome contains two orthologs of *D27*, one of *CCD7*, two of *CCD8* and one of *MAX1^16^* (Supplementary Figs. 1 and 2 and Supplementary Table 1). There are no CYP722 family genes in any of the investigated *Marchantia* species^16, 24^. *CCD8* and *MAX1* orthologs exist in all other AM symbiosis-forming *Marchantia* species tested (Supplementary Figs. 1 and 2). Despite the polymorphisms present in the genome sequences, the overall structure of the genes is well conserved and the amino acid sequences in the conserved domains show high similarities (Fig. 1c). In contrast to this, both *CCD8* and *MAX1* orthologs are absent from the genome of *M. polymorpha*, a non-AM symbiosis-forming species, indicative of gene loss^24^. Together, these phylogenetic analyses suggest that AM symbiosis-forming *Marchantia* species produce SLs.

### Bryosymbiol (BSB), a previously unidentified SL is produced by *M. paleacea*

The correlation between the presence of the *CCD8* and *MAX1* genes and the occurrence of AM symbiosis suggested the possibility that AM symbiosis-forming *Marchantia* species produce SLs. Using LC-MS/MS analysis, we detected carlactonoic acid (retention time of 11.86 min) and its unknown analogue (retention time of 10.73 min) in the rhizoid exudates of *M. paleacea* (Figs. 2a and 2b). Next, to determine whether the carlactonoic acid in *M. paleacea* is synthesized through the same pathway as in flowering plants, we generated knock-out mutants of the two *CCD8* genes (named Mpa*ccd8a* Mpa*ccd8b*, or Mpa*ccd8a/8b*) and one *MAX1* (Mpa*max1*) gene using CRISPR/Cas9 (Supplementary Table 2). Carlactonoic acid was not detected in either Mpa*ccd8a/8b* or Mpa*max1* mutants, indicating that carlactonoic acid biosynthesis is dependent on both MpaCCD8 and MpaMAX1 (Fig. 2a).

**Figure 2.**
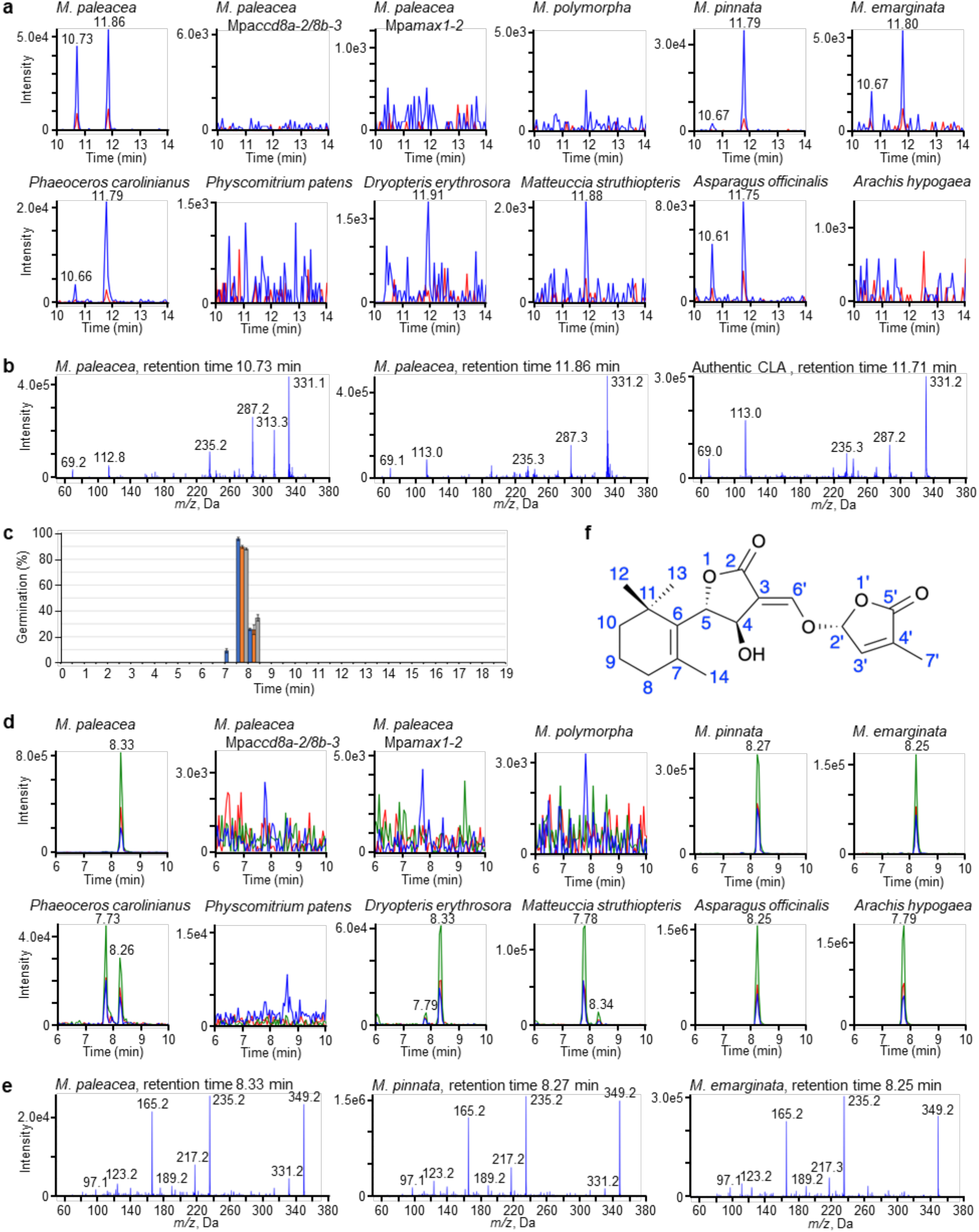
Bryosymbiol (BSB), a previously unidentified SL is produced by *M. paleacea*. (**a**) Detection of carlactonoic acid. Carlactonoic acid in the exudates of different *Marchantia* species and the hornwort *Phaeoceros carolinianus*, protonema extract of the moss *Physcomitrium patens*, root exudates of ferns *Dryopteris erythrosora* and *Matteuccia struthiopteris*, and seed plants *Asparagus officinalis* and *Arachis hypogaea* were analyzed by LC-MS/MS. Multiple reaction monitoring (MRM) chromatograms of carlactonoic acid (blue, 331.10/113.00; red, 331.10/69.00; *m/z* in negative mode) by LC-MS/MS are shown, (**b**) Identification of carlactonoic acid. Product ion spectra derived from the precursor ion (*m/z* 331 in negative mode) of the peaks detected in the exudates of *M. paleacea* and authentic carlactonoic acid (CLA) are shown, (**c**) Germination stimulation activity of the exudate of *M. paleacea* on root parasitic plants. The exudate was separated by reversed-phase HPLC every 30 seconds. All the fractions were tested for seed germination activity on *Orobanche minor* (blue), *Striga hermonthica* (orange) and *Phelipanche ramosa* (gray). Data are means ± SE (approx. 30 seeds per disk, n = 3). (**d**) Detection of BSB. BSB in the exudates of *Marchantia* species and the hornwort *P. carolinianus*,protonema extract of the moss *P. patens*, root exudates of ferns *D. erythrosora* and *M. struthiopteris*, and seed plants *A. officinalis* and *A hypogaea* were analyzed by LC-MS/MS. MRM chromatograms of BSB (blue, 349.00/97.00; red, 349.00/165.00; green, 349.00/235.00; *m/z* in positive mode) are shown, (**e**) Product ion spectra of BSB. Product ion spectra derived from the precursor ion (*m/z* 349 in positive mode) of peaks detected in *Marchantia* species are shown, (**f**) Chemical structure of bryosymbiol (BSB). The absolute stereochemistry at C-4 and C-5 could be interchangeable.

To test for the presence of downstream SL activity, we used a parasitic seed germination assay. This is based on the ability of obligate root parasitic Orobanchaceae plants, such as witchweed (*Striga* spp.) and broomrape (*Orobanche* and *Phelipanche* spp.), to use SLs as host recognition signals in the rhizosphere^25^. This assay is sensitive to concentrations of SLs as low as 10 pM. We fractionated the rhizoid exudates of *M. paleacea* by reversed-phase HPLC and found germination stimulation activity in the fraction collected at around 8 min for all three species of parasitic plants (Fig. 2c). Since there is no known SL eluting at this retention time^26^, we hypothesized that *M. paleacea* produces an unidentified SL. To identify the nature of this compound, we collected and purified the active fraction from the rhizoid exudate of *M. paleacea* in a large-scale culture. The molecular formula of the purified amorphous solid (0.33 mg) was established to be C_19_H_24_O_6_ based on a protonated ion obtained by HR-ESI-TOF-MS, and its structure was determined by NMR spectroscopy, ESI- and EI-MS spectrometry (Figs. 2d and 2e, Supplementary Figs. 3a-c and Supplementary Table 3). The chemical structure of the identified SL was determined as (*E*)-5-((4-hydroxy-2-oxo-5-(2,6,6-trimethylcyclohex-1-en-1-yl)dihydrofuran-3(2*H*)-ylidene)methoxy)-3-methylfuran-2(5*H*)- one (Fig. 2f). The CD spectrum indicated that it has a 2′(*R*) configuration like previously identified natural SLs (Supplementary Fig. 3d). We named this newly identified SL bryosymbiol, or BSB. Although further studies are needed to confirm the absolute stereochemistry of BSB, the NOESY correlations suggested the relative stereochemistry at C-4 and C-5 is *anti* and we tentatively assigned BSB to be (4*R*, 5*S*, 2′*R*) (Supplementary Fig. 3a and Supplementary Table 3).

BSB was detected in the exudate and plant extracts of WT *M. paleacea* but not in the Mpa*ccd8a/8b* and Mpa*max1* mutants (Figs. 2d and 3a). In contrast, we detected a higher level of carlactone accumulation in the extract of the Mpa*max1* mutant than that of the WT (Fig. 3b). This suggests that the substrate of MpaMAX1 is likely to be carlactone, similar to that of MAX1 homologs in vascular plants. To determine whether MpaMAX1 directly converts carlactone to BSB, we performed *in vitro* metabolic experiments. When the microsomal fraction of yeast expressing Mpa*MAX1* was incubated with carlactone, the production of carlactonoic acid and BSB were detected (Fig. 3c). This indicates that MpaMAX1 catalyzes two steps, namely the conversion from carlactone to carlactonoic acid and carlactonoic acid to BSB. To further confirm this pathway, we incubated microsomes of yeast expressing *MpaMAX1* with carlactonoic acid and detected BSB (Fig. 3c). We concluded that MpaMAX1 produces BSB from carlactone via carlactonoic acid. The conversion of carlactonoic acid to BSB is presumed to be via epoxidation and that upon proton abstraction of the carboxyl group, closure of the C ring occurs (Fig. 3d). Stereoselective closure of the C ring may occur, producing BSB (4*R*,5*S*,2′*R*) and its diastereomer (4*S*,5*R*,2′*R*), probably depending on the epoxy configuration. However, the occurrence of 7,8-epoxy-carlactonoic acid, a putative intermediate between carlactonoic acid and BSB (Fig. 3d), could not be confirmed due to the lack of the authentic standard.

**Figure 3.**
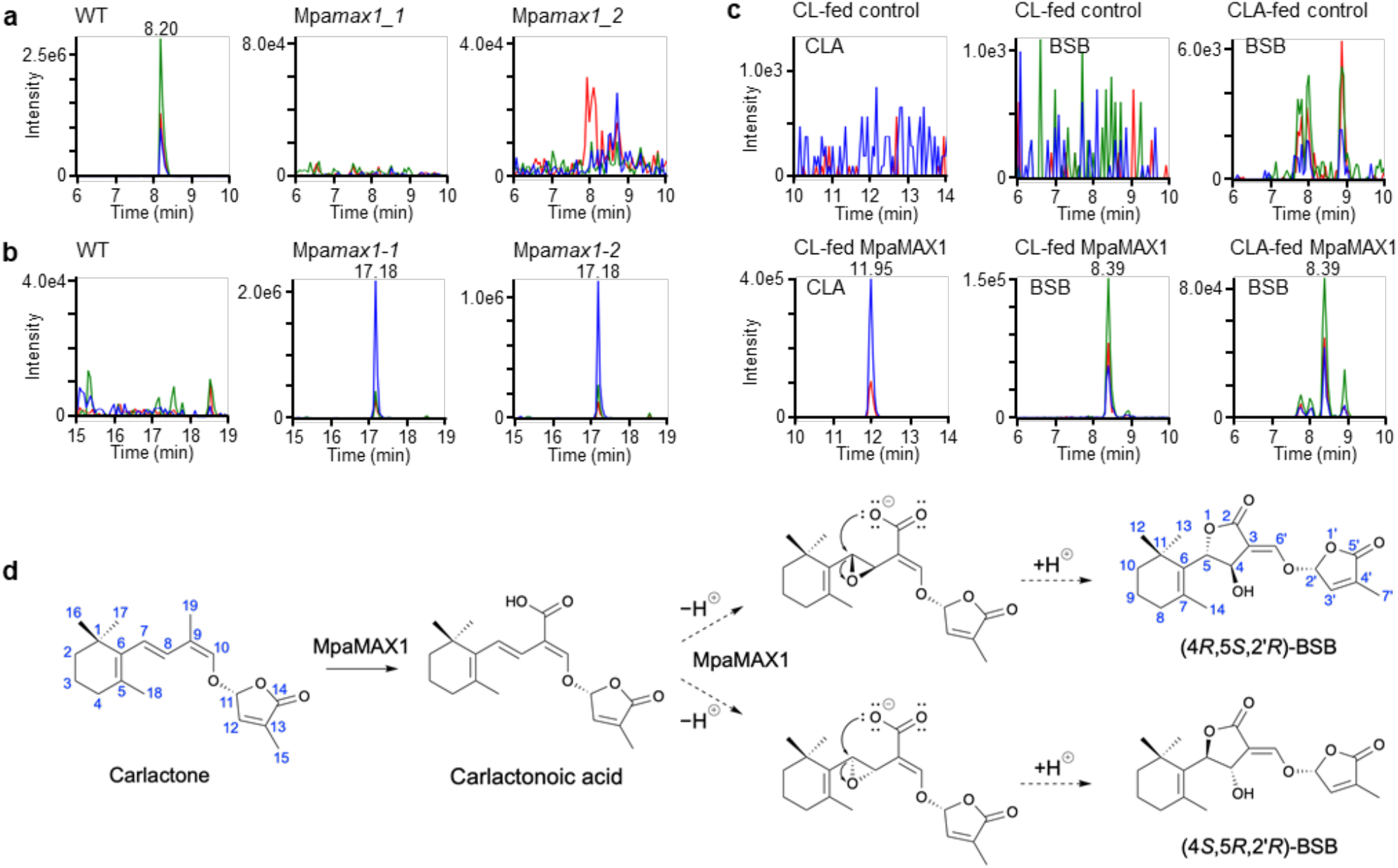
MpaMAX1 produces BSB from carlactone via carlactonoic acid. **(a and b)** Detection of BSB and carlactone in plant tissues of *Marchantia* Whole plants of WT and the Mpa*max1* mutant (two alleles) were extracted and analyzed by LC-MS/MS. MRM chromatograms of BSB (a) and carlactone (b) (blue, 303.00/97.00; red, 303.00/189.00; green, 303.00/207.00; *m/z* in positive mode) are shown (**c**) Detection of carlactonoic acid and BSB in feeding experiments with recombinant MpaMAX1. *Rac*-carlactone (CL) or rac-carlactonoic acid (CLA) was incubated with the microsomes of MpaMAX1-expressed yeast. Yeast microsomes with an empty vector were used as a control. MRM chromatograms of carlactonoic acid and BSB in extracts of the microsomes are shown, (**d**) Proposed mechanism of the conversion to BSB from carlactone. MpaMAX1 oxidizes C-19 to produce carlactonoic acid and catalyzes the α-or β-epoxidation of Δ7,8. Upon proton abstraction of the carboxyl group, the stereoselective closure of the C ring occurs, producing BSB isomers depending on the epoxy configuration.

To determine whether BSB is a derived trait of *M. paleacea* or more widespread across the plant diversity, we investigated its presence in diverse land plants. First, we analyzed *M. pinnata, M. emarginata* and *M. polymorpha*. Carlactonoic acid and BSB were detected in exudates from the two AM symbiosis-forming *Marchantia* species but not from *M. polymorpha*, consistent with the presence/absence of *CCD8* and *MAX1* (Figs. 2a, 2d and 2e and Supplementary Figs. 1 and 2). Neither carlactonoic acid nor BSB was detected in the extracts of the model moss *Physcomitrium patens*, which lacks the *MAX1* gene (Figs. 2a and 2d)^27^. Since *CCD8* and *MAX1* are present in hornworts^28, 29^, we analyzed exudates from the hornwort *Phaeoceros carolinianus*. We also analyzed several ferns and seed plants. Carlactonoic acid and BSB (retention time of 8.3 min), and/or a likely stereoisomer of BSB (retention time of 7.7 min), were detected in exudates from the hornwort *P. carolinianus*, the ferns *Dryopteris erythrosora* and *Matteuccia struthiopteris*, and the seed plants *Asparagus officinalis* (monocot) and *Arachis hypogaea* (dicot) (Figs. 2a and 2d and Supplementary Fig. 4).

Collectively, these results demonstrate that BSB is produced by AM symbiosis-forming *Marchantia* and a variety of vascular plants. Because bryophytes, such as liverworts and hornworts, and vascular plants, such as ferns and flowering plants, diverged more than 400 million years ago, it is inferred that BSB was likely present in the most recent common ancestor of land plants.

### Enhanced BSB synthesis correlates with an AM symbiosis permissive state

In flowering plants, phosphate starvation stimulates the expression of SL-biosynthesis genes and the actual production of SLs, and correlates with a permissive state for the AM symbiosis to be established^7, 30^. We tested whether BSB synthesis is also enhanced in *M. paleacea* under phosphate-deficient conditions. Expression of all five genes in the SL synthesis pathway, namely, Mpa*D27*, Mpa*CCD7*, Mpa*CCD8A*, Mpa*CCD8B* and Mpa*MAX1*, were increased upon phosphate starvation (Fig. 4a). We also found that the expression of the *CCD8* gene in *Anthoceros agrestis*, a hornwort, was induced by phosphate starvation (Fig. 4a).

**Figure 4.**
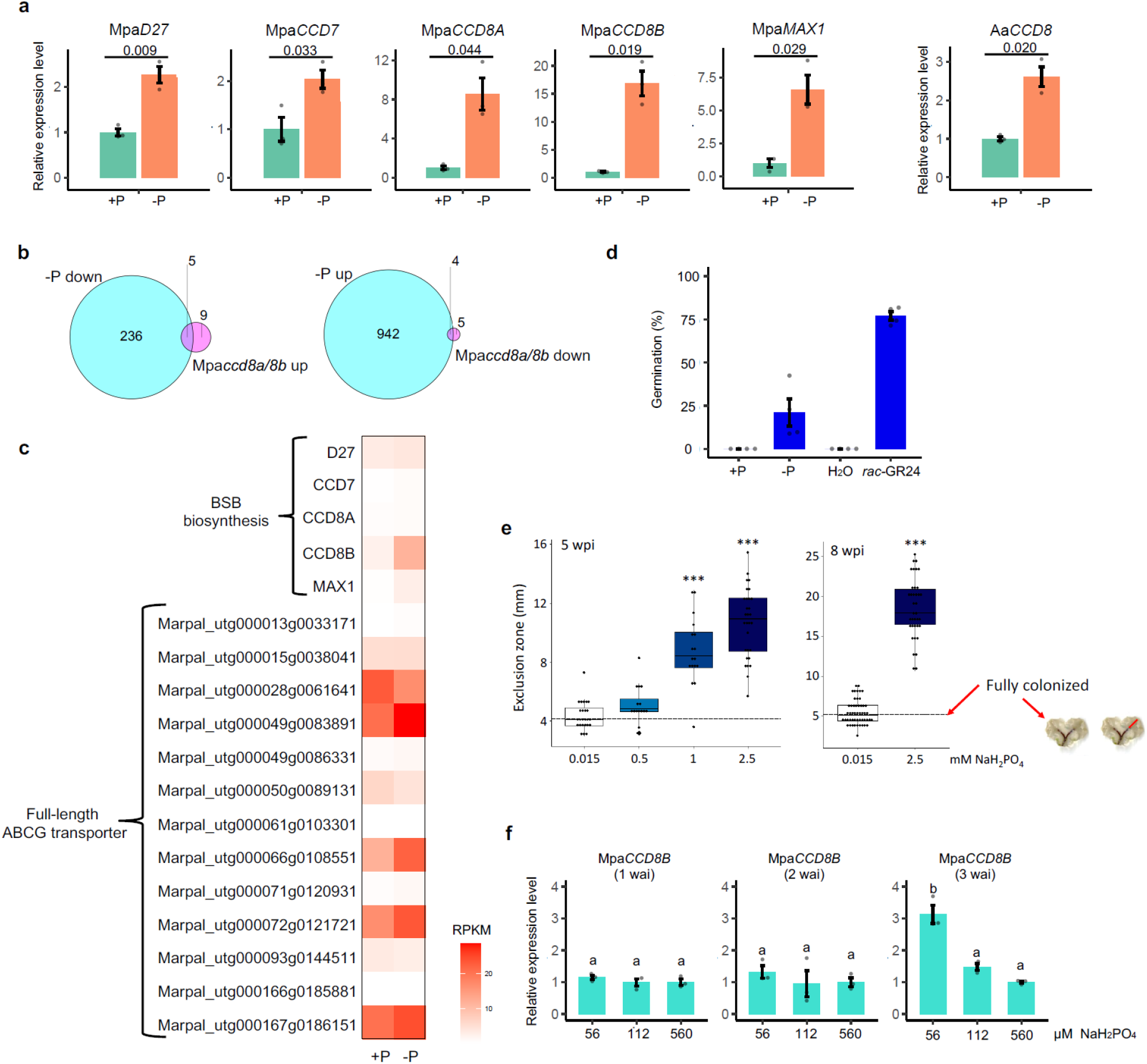
Enhanced BSB synthesis correlates with an AM symbiosis permissive state. **(a)** Increased expression of the SL biosynthesis genes, Mpa*D27*, Mpa*CCD7*, Mpa*CCD8A* Mpa*CCD8B* and Mpa*MAX1*, of *M. paleacea*, and AaCCDS of *Anthoceros agrestis* under phosphate starvation. Plants were grown on half-strength Gamborg’s B5 medium with (+P) or without (-P) phosphate, **(b)** Identification of DEGs under phosphate starvation and in Mpa*ccd8a/8b* mutants by RNAseq analysis. Venn diagrams of genes down-regulated in -P condition compared to +P condition and up-regulated in Mpa*ccd8a/8b* compared to WT (left) and up-regulated in -P condition compared to +P condition and down-regulated in Mpaccd8a/8b compared to WT (right), **(c)** Heatmap of gene expression under phosphate deficiency in *M. paleacea*. **(d)** Increase in the stimulation activity of the exudate of *M. paleacea* on seed germination of *Orobanche minor* under phosphate starvation. Exudates from *M. paleacea*grown on +P and -P medium were used for analysis, (**e**) Low phosphate allows mycorrhizal colonization. Plants inoculated with *Rhizophagus irregularis* were watered with Long Ashton solution containing increasing phosphate concentration and the distance (mm) between the notch and the colonized area of the thalli was measured (n ≤ 15). Higher distance signifies arrest of the colonization. Stars show statistical significance(ANOVA, post hoc Tukey). **(f)** Developmental stage-dependent induction of Mpa*CCD8B* expression in response to low phosphate conditions. Gemmae of 1-3 weeks after inoculation (wai) were transferred to media containing different concentrations of phosphate. Values in (a) and (d) are means ±SD (n = 3, n=4, respectably), and significant differences detected with Welch’s t-test are indicated (a and d). Values in (f) are means ±SD (n =3) and The HSD test was used for multiple comparisons. Statistical differences (P-values < 0.05) are indicated by different letters.

To obtain a genome-wide view of the phosphate level-dependent gene expression in *M. paleacea*, we performed RNAseq analysis (Fig. 4b). The expression of many genes changed under phosphate-deficient conditions (Supplementary Table 4). In petunia and *Medicago truncatula*, expression of *PDR1*, an SL transporter gene, is also enhanced under low phosphate conditions, implying that increased amounts of SLs are exuded to the rhizosphere under phosphate starvation^31, 32^. Although SL transporters in bryophytes have not yet been identified, ABCG transporter genes, homologs of *PDR1*, were up-regulated under phosphate-deficient conditions in *M. paleacea* (Fig. 4c).

We then tested the level of exuded BSB using the parasitic seed germination assay under low phosphate conditions. Consistent with the enhanced expression of SL biosynthesis and putative transporter genes, seed germination frequency was extremely higher in exudates of *M. paleacea* grown under phosphate-deficient conditions (Fig. 4d), implying that the amount of BSB exuded was increased upon phosphate starvation. We finally examined if phosphate starvation affects AM symbiosis in *M. paleacea* as in vascular plants (Fig. 4e). While phosphate-deficient conditions were permissive for AM symbiosis, increasing level of phosphate in the watering solution resulted in a gradual decrease in colonization by the AM fungus, illustrated by an increased zone without AM symbiosis (Fig. 4e).

AM symbiosis does not occur during the early stages of *M. paleacea* development and is first observed in three-week-old gemmae. Thus, we also tested the developmental control of BSB synthesis. We found enhanced *MpaCCD8B* expression under low phosphate conditions in three-week-old gemmae but not in earlier stage gemmae (Fig. 4f). Altogether, these results indicate that BSB synthesis correlates with a permissive state for AM symbiosis in *M. paleacea*.

### BSB is required for symbiosis with AM fungi in *M. paleacea*

The correlation between the presence of BSB and AM symbiosis suggests that bryophytes use BSB as a rhizosphere signaling chemical to promote the establishment of symbiosis with AM fungi. We confirmed that BSB induces hyphal branching in the AM fungus *Gigaspora margarita* with a minimum effective concentration of 10 pg/disk, which is within the range of the effects of previously described natural SLs (1 to 100 pg/disk) (Fig. 5a)^33^. To further test the symbiotic function of BSB *in vivo*, the Mpa*ccd8a*, Mpa*ccd8b* and Mpa*ccd8a/8b* mutants were inoculated with spores of the AM fungus *Rhizophagus irregularis*. Six weeks after inoculation, the Mpa*ccd8a* and Mpa*ccd8b* mutant thalli showed colonization similar to the WT. In contrast, almost all thalli of the Mpa*ccd8a/8b* mutants were not colonized (Figs. 5b and 5c and Supplementary Fig. 5). To determine which stages of the symbiotic establishment were affected in the mutants, the number of infection points was scored. While the number of infection points was similar in the WT plants and the Mpa*ccd8a* and Mpa*ccd8b* single mutants, all three alleles of the Mpa*ccd8a/8b* mutant showed no or very few infection sites (Figs. 5c and Supplementary Fig. 5). To further determine whether the lack of infection was due to the absence of stimulation of hyphal branching by BSB, complementation assays were conducted using *rac*-GR24, a synthetic analogue of SL known to induce hyphal branching. Exogenous treatment with 100 nM *rac*-GR24 restored colonization and the number of infection points in the three Mpa*ccd8a/8b* mutant alleles (Figs. 5b and 5c and Supplementary Fig. 5).

**Figure 5.**
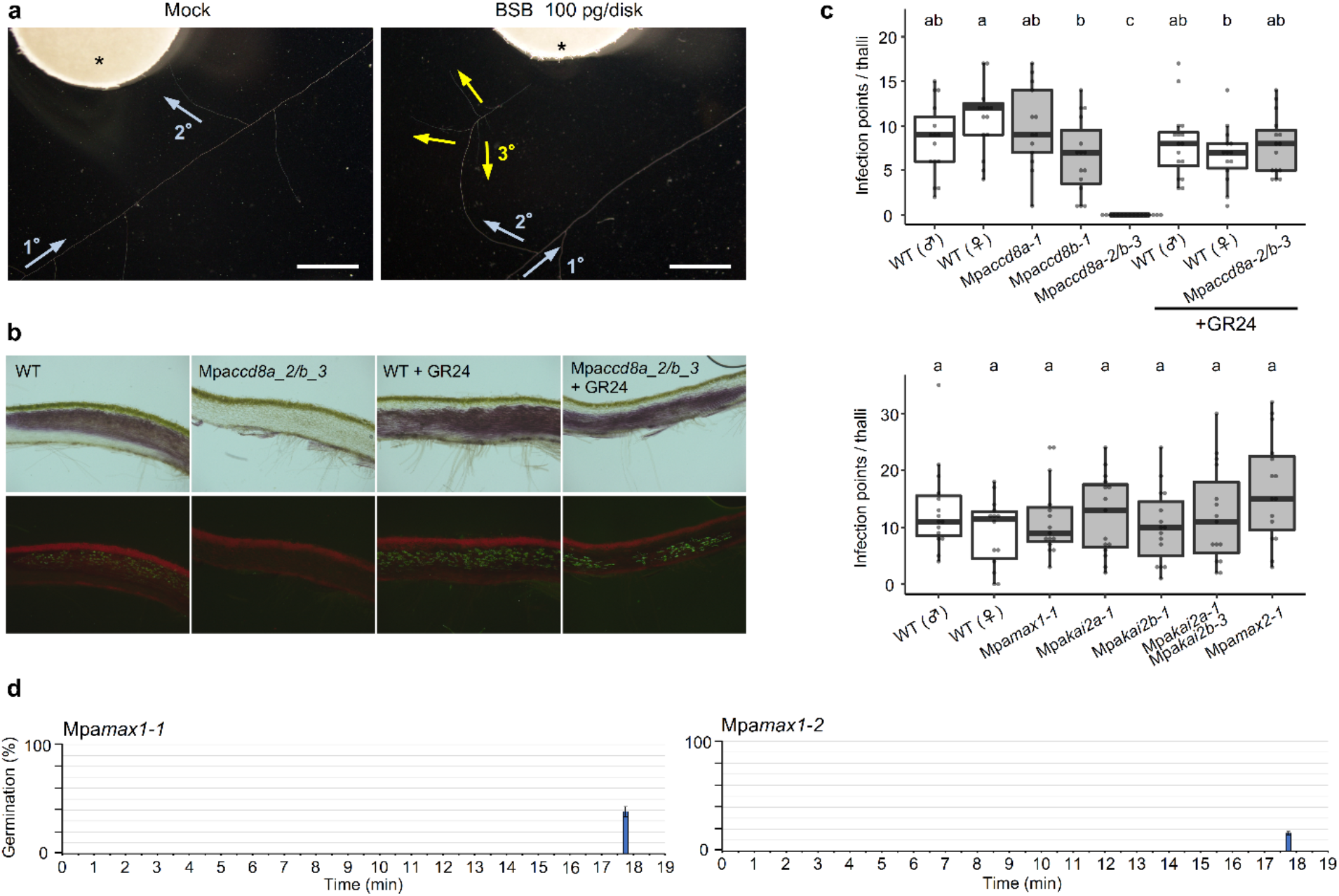
BSB is required for symbiosis with AM fungi in *M. paleacea*. (**a**) Hyphal branching of *Gigaspora margarita* induced by BSB (100 pg per disk) using the paper disk diffusion method. Arrows indicate the direction of growth of primary, secondary and tertiary hyphae. Asterisks indicate paper disks, (**b**) Upper and bottom panels are transverse sections of colonization by *Rhizophagus irregularis*. The bottom panels show WGA-FITC staining of the fungi. Scale bars: 0.5 mm. (**c**) The number of infection points per thalli (n ≥ 13). Letters show different statistical groups (ANOVA, post hoc Tukey). (**d**) Germination stimulation activity on root parasitic plants of the exudates of the Mpamaxf mutant. The exudates of the two alleles were separated by reversed-phase HPLC. Every 30 sec all the fractions were tested for seed germination activity on *Orobanche minor*. Data are means ± SE (approx. 30 seeds per disk, n =3).

In flowering plants, D14LIKE (D14L), an ortholog of KARRIKIN INSENSITIVE2 (KAI2), and D3/MAX2-dependent signaling positively regulate AM symbiosis^34, 35^. In contrast to this, AM symbiosis was not affected in *kai2* or *max2* mutants of *M. paleacea* (Fig. 5c and Supplementary Fig. 5). This suggests that KAI2-dependent signaling was recruited to control AM symbiosis in vascular plant lineages.

In contrast to the Mpa*ccd8a/8b* mutants, neither colonization nor infection sites were affected in *Mpamax1* mutants, despite the absence of BSB production (Fig. 5c and Supplementary Fig. 5). To understand this discrepancy, we pursued the presence of an active SL other than BSB in the exudates of Mpa*max1* mutants. We fractionated the exudates from Mpa*max1* by reversed-phase HPLC and tested for the presence of SL-like activity using the parasitic seed germination assay. These assays identified a single active fraction at the retention time of 18 min (Fig. 5d), while there was no activity in the corresponding fraction of WT exudates (Fig. 2c). This fraction contains carlactone (Fig. 3b), which is known to induce hyphal branching, although at higher concentrations than BSB^36^. Accumulation of carlactone in the Mpa*max1* mutants may explain the observed normal fungal colonization.

From this reverse genetic approach and chemical complementation assays, we conclude that the function of SLs as symbiotic signal in the rhizosphere is an ancestral trait in land plants, which has been maintained in both the vascular plants and the bryophytes for 400 million years.

### BSB-deficient mutants of *M. paleacea* show no abnormal developmental patterns and gene expression profiles

We next examined the possible endogenous functions of BSB in *M. paleacea*. Despite the absence of BSB synthesis, no alteration in growth pattern or morphology was observed in Mpa*ccd8a/8b* and Mpa*max1* loss-of-function mutants throughout their growth, even in plants grown under the low phosphate conditions that stimulate BSB synthesis (Figs. 6a-d). This is consistent with the results of the RNAseq analysis comparing gene expression between WT and Mpa*ccd8a/8b*, which shows that only a small number of genes are differentially regulated (Fig. 4b). These data suggest that the absence of endogenous BSB causes little effect on *M. paleacea*. Feedback regulation of SL synthesis genes such as *CCD8* is observed in many seed plants^7, 37^, whereas expression of Mpa*CCD8A* and Mpa*CCD8B* genes are not affected by exogenously applied SLs (Fig. 6e).

**Figure 6.**
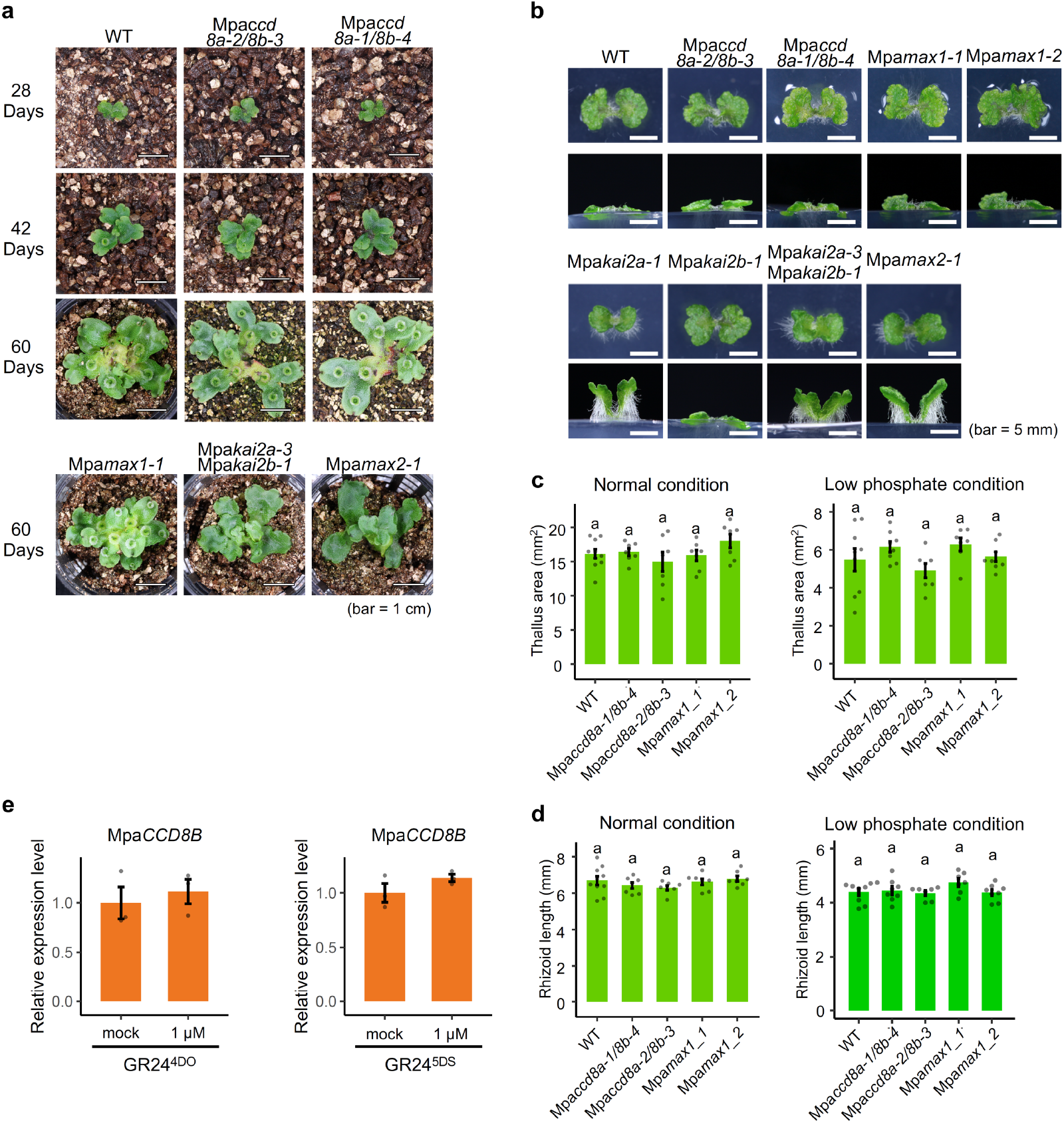
BSB-deficient mutants of *M. paleacea* show no abnormal developmental patterns and gene expression profiles. (a and b) Growth of BSB-deficient (Mpa*ccd8a-2/8b-3, Mpaccd8a-1/8b-4* and Mpa*max1-1* and Mpa*mxf-2*) and KAI2 signaling (Mpa*kai2a-1, Mpakai2b-1*, Mp*aka/2a-3* Mpa*kai2b-1* and Mpa*max2-1*) mutants of *M. paleacea* grown in soil for 28, 42 and 60 days (a) and in media for 20 days (b). (c and d) Thallus area and rhizoid length of BSB-deficient mutants of *M. paleacea* grown on half-strength Gamborg’s B5 medium containing 560 μM (c) and 56 μM (d) phosphorus (NaH_2_PO_4_) for 14 days, (e) Expression of Mpa*CCD8B* in WT plants after GR24^4D^ and GR24^5DS^ treatments. Values in (c) and (d) are means ± SD (n ≧ 7) and in (e) are means +SD (n = 3). The HSD test was used for multiple comparisons and statistical differences (P-values < 0.05) are indicated by different letters.

In flowering plants, SL is perceived by D14, encoding an α/β-hydrolase protein^38, 39, 40^. *D14* originated from a gene duplication of *KAI2* before the radiation of seed plants^41^. Although endogenous ligands of KAI2 are unknown, it is thought that D14 and KAI2 interact with other signaling components such as the F-box protein MAX2/D3 upon the perception of their cognate ligands^8^. This leads to the degradation of SMXL proteins that work as regulators of transcription of downstream genes^42, 43^. We previously reported that KAI2, MAX2/D3 and SMXL function in a common proteolysis-dependent signaling pathway in *M. polymorph*a^44^. Similarly to *M. polymorpha*, the *M. paleacea* genome contains one MAX2/D3 (Mpa*MAX2*) and two KAI2-like genes (*MpaKAI2A* and *MpaKAI2B*) but no *bona fide D14*, as is the case in other nonseed plants. To pursue a possibility that BSB plays an endogenous function in *M. paleacea*, we tested the possibility that MpaKAI2A and/or MpaKAI2B act as the receptors of BSB. *Mpakai2a* and *Mpamax2* mutants (Supplementary Table 2) showed altered growth patterns, including reduced and upward growth of the early-stage thalli and delayed gemmae cup initiation (Figs. 6a and 6b). Such developmental defects were not observed in the Mpa*ccd8a/8b* or the Mpa*max1* mutants (Figs. 6a and 6b). To further investigate the potential link between MpaKAI2A and MpaKAI2B and SL, their physical interaction with four stereoisomers of the synthetic SL GR24 was investigated by differential scanning fluorimetry (DSF) analysis (Supplementary Figs. 6a and 6b). Only GR24^ent-5DS^, which has an unnatural D-ring configuration and has been shown to interact with Arabidopsis KAI2^45^, was able to weakly interact with MpaKAI2A whereas no sign of interaction was observed between MpaKAI2B and GR24. In flowering plants, SL synthesis is enhanced in SL signaling mutants because of the loss of feedback regulation^46^. However, no such increase was observed in the levels of carlactonoic acid and BSB in exudates of Mpa*kai2a/2b* and Mpa*max2* mutants (Supplementary Figs. 6c-f). Altogether, these phenotypic and biochemical analyses indicate that MpaKAI2A, MpaKAI2B and MpaMAX2 are not involved in the perception of BSB or other metabolites downstream CCD8 or MAX1 in *M. paleacea*.

### Heterologous expression of AtD14, the SL receptor of Arabidopsis, renders *M. paleacea* cells BSB-sensitive

From the previous phylogenetic studies and the experimental work presented above, it can be proposed that BSB does not function as a hormone in bryophytes due to the absence of cognate receptors. In other words, although an SL, BSB, is produced and the known signaling pathway is present in bryophytes, the lack of the upstream receptor results in the two pathways being disconnected. If this hypothesis is valid, mimicking the seed-plant duplication of the KAI2-like gene by adding a cognate SL receptor in *M. polymorpha* and *M. paleacea* should lead to SL responsiveness. To experimentally test this hypothesis, we took a gain-of-function approach in which the canonical SL receptor D14 of Arabidopsis (*AtD14*) was fused with the promoter of *M. polymorpha ELONGATION FACTOR1* (*EF1*) gene and introduced into *M. polymorpha* and *M. paleacea*. We examined whether endogenous BSB is recognized by the introduced AtD14, leading to signal transduction. More than 20 independent lines in each species were produced and two lines that show high and low levels of At*D14* expression were selected from each species (named Mp*AtD14*ox and Mpa*AtD14*ox, for M*. polymorpha* and *M. paleacea*, respectively) for further analysis (Fig. 7a and Supplementary Fig. 7a). First, we tested if the introduction of AtD14 enables SL perception and signal transduction in response to *rac*-GR24 in *M. polymorpha* cells that do not produce BSB. The *DIENELACTONE HYDROLASE LIKE PROTEIN1* (*DLP1*) and *MpSMXL* genes of *M. polymorpha*, whose expression is highly dependent on KAI2-dependent signaling, were used as markers^44^. Expression of *MpDLP1* and *MpSMXL* was induced by the application of *rac*-GR24 in the *MpAtD14*ox lines but not in WT plants (Supplementary Fig. 7b). To verify that AtD14 functions in the same signaling pathway as the endogenous KAI2A, Mp*kai2a* mutation was introduced into the Mp*AtD14*ox lines. By contrast with the Mp*kai2a* mutants, the upward thallus growth phenotype was rescued by application of GR24^4DO^ and GR24^5DS^, which have a 2’(*R*) configuration as natural SLs^45^, in the MpAt*D14*ox/Mp*kai2a* lines (Supplementary Fig. 7c).

**Figure 7.**
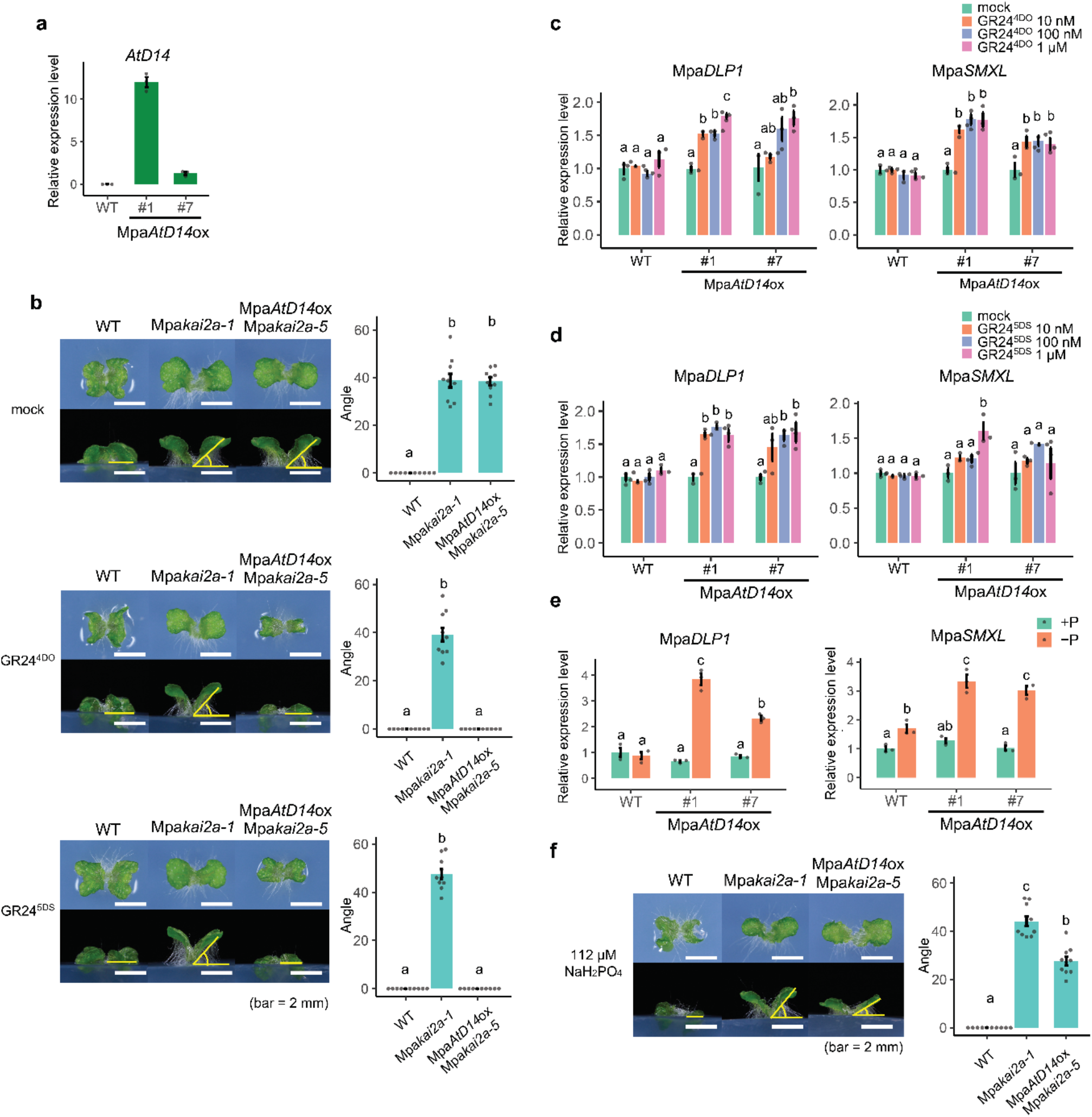
Heterologous expression of *AtD14*, the SL receptor of Arabidopsis, renders *M. paleacea* cells BSB-responsive. **(a)** Expression of *AtD14* introduced into *M. paleacea*. **(b)** Complementation of Mpa*kai2a* phenotypes in Mpa*AtD14*ox/Mpa*kai2a* (*M. paleacea*) lines by addition of 1 μM GR24^4DO^ or 1 μM GR24^5DS^. **(c and d)** Induction of Mpa*DLP1* and Mpa*SMXL*, marker genes of KAI2-dependent signaling, in response to SL treatment. Expression of Mpa*DLP1* and Mpa*SMXL* in WT and Mpa*AtD14*ox lines in *M. paleacea* after GR24^4DO^ (c) or GR24^5DS^ (d) treatment are shown, (e) Induction of Mpa*DLP7* and Mpa*SMXL* expression by phosphate starvation in Mpa*AtD14*ox lines of *M. paleacea*. **(f)** Complementation of Mpa*kai2a* phenotypes in Mpa*AtD14*ox/Mpa*kai2a* (*M. paleacea*) lines grown on media with a reduced phosphorus level. Values in (a) and (c) to (e) are means ± SD (n = 3), and values in (b) and (f) are means ± SD (n = 10). The HSD test was used for multiple comparisons. Statistical differences (P-values < 0.05) are indicated by different letters.

Phenotypes of Mpa*kai2a*, the loss-of-function mutant of *KAI2A* in *M. paleacea*, closely resembled those of Mp*kai2a* (Figs. 6b and 7b). As for *M. polymorpha*, the thallus upward curving was rescued by application of GR24^4DO^ and GR24^5DS^ in the Mpa*AtD14*ox/Mpa*kai2a* (Fig. 7b). These results indicate that introducing AtD14 enables SLs perception and downstream signal transduction through the KAI2-dependent signaling pathway in both *M. paleacea* and *M. polymorpha*.

We next directly tested the effect of exogenously applied SLs on the signaling cascade. As is the case in *M. polymorpha*, neither GR24^4DO^ nor GR24^5DS^ enhanced expression of the Mpa*DLP1* and Mpa*SMXL* genes in WT *M. paleacea* plants, even at the highest concentration (Figs. 7c and 7d). This implies that the applied SLs are not perceived by WT *M. paleacea* cells, consistent with the disconnection between SL and the KAI2-like signaling pathway. In contrast, in the Mpa*AtD14*ox lines containing the SL receptor gene, expression of Mpa*DLP1* and Mpa*SMXL* was enhanced by the application of either GR24^4DO^ or GR24^5DS^ in a concentration-dependent manner (Figs. 7c and 7d).

We showed that BSB synthesis is enhanced under phosphate starvation (Figs. 4a and 4c). Therefore, we hypothesized that enhanced synthesis of BSB may recruit AtD14-dependent signaling under phosphate-deficient conditions. Indeed, expression of Mpa*DLP1* and Mpa*SMXL* was induced in Mpa*AtD14*ox lines under phosphate starvation (Fig. 7e). Although a slight increase in Mpa*SMXL* expression under phosphate depletion was sometimes observed in WT plants, the same trend was detected in Mpa*ccd8a/8b*, suggesting that the slight increase was caused by unknown SL-independent mechanisms (Supplementary Fig. 7d). Besides the activation of marker genes, we tested whether the gain of the cognate SL receptor was sufficient to recruit BSB as an endogenous signal with a physiological effect. Thus, Mpa*kai2a* and Mpa*AtD14*ox/Mpa*kai2a* lines were transferred to a medium with a lower level of phosphate. On this medium, while the upward curving phenotype of the Mpa*kai2a* lines remained fully unaffected, this defect was partially, but significantly, rescued in the Mpa*AtD14*ox/Mpa*kai2a* lines (Fig. 7f). Taken together, these results indicate that endogenous BSB can be perceived by a newly integrated SL receptor and transduce the signal through the MpaMAX2-dependent signaling pathway; that is, BSB is perceived in *M. paleacea* cells when the cognate receptor is present. This strongly supports the hypothesis that the evolution of SLs as an endogenous hormone is only dependent on the evolution of a cognate receptor in the KAI2 clade.

## DISCUSSION

### The ancestral function of SLs is as symbiotic signaling molecules in the rhizosphere

In flowering plants, SLs play dual roles as a class of phytohormones to control various aspects of growth and development and as rhizosphere signaling chemicals to control symbiosis with AM fungi and parasitism by root parasitic plants^5, 6, 7, 47^. AM symbiosis is found in most extant land plants, which comprise two main monophyletic clades, the non-vascular bryophytes and the vascular tracheophytes and was present in the last common ancestor of land plants more than 400 million years ago^48^. In this study, we showed that the SL, BSB, is produced in a wide range of plant species, including flowering plants, indicating that BSB is an ancestral and conserved form of SLs. In *M. paleacea*, BSB acts as a rhizosphere signaling molecule essential for symbiosis with AM fungi. Thus, it can be concluded that the function of SLs as signaling molecules to communicate with AM fungi was already established in the common ancestor of the bryophytes and vascular plants.

AM symbiosis was an innovation by the first land plants to acquire minerals, in particular phosphate, from the nutrient-poor soil they faced during the colonization of emerged land. In all mutualistic systems, control mechanisms have evolved on both the host and symbiont sides to optimize the benefits of the symbiosis^49^. In flowering plants, SL synthesis is regulated by phosphate availability^30, 50, 51^. Upon phosphate starvation, SL biosynthesis and transporter genes are enhanced resulting in an increase in the level of SLs exuded to the rhizosphere^32^. This leads to enhanced AM symbiosis and consequently, to increased phosphate supply. We showed that phosphate-dependent control of SL synthesis occurs in the liverwort *M. paleacea* and the hornworts *Anthoceros agrestis*. These results imply that the control of AM symbiosis through phosphate-dependent SL synthesis was already present in the most common ancestors of bryophytes and vascular plants.

### Convergent recruitment of SL as hormonal signal

Although we cannot rule out that BSB does play an endogenous role in *M. paleacea*, the absence of observed growth phenotypes in the Mpa*max1* and Mpa*ccd8a/8b* mutant, and the loss of both *CCD8* and *MAX1* in *M. polymorpha*, a non-symbiotic species, suggest that its function is exclusively symbiotic in liverworts. Even more certain is the disconnection between BSB and the KAI2/MAX2 signaling pathway. Contrary to the loss of *CCD8* and *MAX1* in *M. polymorpha, KAI2A, KAI2B* and *MAX2* are conserved in *M. polymorpha* despite its non-symbiotic characteristics and the developmental phenotypes observed in *M. paleacea kai2a* and *max2* mutants resemble the ones observed in *M. polymorpha* but the phenotypes are absent in Mpa*ccd8a/8b* or Mpa*max1* mutant. Loss of *CCD8* is also observed in the water ferns *Azolla filiculoides* and *Salvinia cucullata* that have also lost AM symbiosis^52^. Such a correlation between gene and trait losses is indicative of a specific function of the genes, here in AM symbiosis. Thus, it can be inferred that the metabolites from the CCD8/MAX1 SL biosynthesis pathway and KAI2/MAX2 signaling were originally independent. The role of the CCD8-derived metabolites as endogenous plant hormones evolved during seed plant diversification by connecting them to receptors from the KAI2/D14 family. Extensive phylogenetic analysis revealed that D14 evolved through putative neo-functionalization following gene duplications of the KAI2 family members in the common ancestor of seed plants, leading to its function as cognate receptor for SLs as plant hormones regulating multiple developmental traits in flowering plants^13, 41, 47^. In this study, we mimicked the evolution of such a hormonal system by introducing the *D14* from Arabidopsis in *M. paleacea* and showed that endogenous BSB and exogenously supplied SL analogs are perceived and trigger signal transduction in these engineered lines through D14-MAX2 dependent signaling.

Despite the lack of *MAX1*, molecules with an SL function are synthesized through CCD8 in *P. patens*^15^. CCD8-derived metabolites have also been recruited for quorum sensing-like functions in *P. patens*^14^. These metabolites are currently elusive but the absence of *MAX1* in *P. patens* suggests that those are neither carlactonoic acid nor BSB. Interestingly, the *KAI2* genes are highly duplicated in *P. patens* and some of them are involved in the perception of the unknown CCD8-derived metabolites, but their signaling is MAX2-independent^53, 54^. These findings imply that *P. patens* generated both the metabolic pathway and the signaling system independently from other bryophytes after the loss of the *MAX1* gene. Similar acquisition of signaling systems occurred in root parasitic plants, in which the *KAI2* genes duplicated massively^47^. Some of the *KAI2* paralogs were identified as extremely sensitive SL receptors that recognize SLs from host plants^47, 55, 56^. Therefore, the generation of SL receptors from KAI2 and of molecules with an SL function occurred repeatedly and independently during the evolution of land plants.

### A model for the evolution of hormonal networks in plants

The origin of the complex hormonal pathways observed in extant flowering plants has been extensively studied by combining reverse genetics conducted in bryophytes and phylogenetic studies on the entire green lineage^57, 58^. From our work and previous studies, we propose a general model for the evolution of hormonal networks in plants. In a first step, the signaling and biosynthesis pathways evolved independently, following a *ligand first* or a *signaling first* modes of evolution. The *ligand first* mode is captured by the hormone auxin and ABA which are produced by green algae whereas the signaling pathways are restricted to land plants, and highly conserved^59, 60, 61^. The *signaling first* mode has been observed for gibberellin (GA), an essential hormone in vascular plants which is absent from bryophytes whereas the central transcriptional regulator involved in the GA pathway, DELLA, is present in bryophytes where it regulates plant growth and stress responses^62, 63^. With both a biosynthetic pathway and a functional signaling pathway present, we propose that the evolution of a receptor is sufficient to evolve a fully functional hormonal network. In the Zygnematophyceae, which are the closest algal clade relative to land plants, the ortholog of the ABA receptor functions independently of ABA^64, 65^. The ABA-receptor ortholog in land plants acquired the ability to bind ABA which was sufficient to recruit its downstream signaling pathway^24, 66^. Here, we revealed that SL evolved following a *ligand first* mode, and experimentally demonstrated that the gain of a receptor was sufficient to recruit SL as hormonal signal. Why such connections between the product of an existing biosynthetic pathway and a signaling pathway emerge and are selected remains an open question. An attractive view is that SLs being exuded under phosphate deprivation conditions may have been first recruited as a rhizospheric signal in plant communities and later as an endogenous mobile signal. Stepwise recruitment of non-connected biosynthetic and signaling pathways into integrated processes might represent a general mechanism for the evolution of intercellular and inter-organism communication.

## Methods

### Plant materials and growth conditions

*Marchantia paleacea* ssp. *diptera*, *M. polymorpha*, *M. pinnata*, *M. emarginata* and *Phaeoceros carolinianus* were harvested from the wild in Japan (*Marchantia paleacea* ssp. *diptera* was collected in Nara prefecture, the others were collected in Hiroshima prefecture). *Dryopteris erythrosora* and *Matteuccia struthiopteris* plants were purchased from a garden store. Seeds of *Asparagus officinalis* (cv. Shower, Takii, Japan) and *Arachis hypogaea* (cv. Chibahandachi, Watanabe Noji, Japan) were used. *M. polymorpha* and *M. paleacea* were grown at 22 °C under continuous light on 1/2 Gamborg’s B5 medium (pH 5.5) with 1 % agar (Guaranteed Reagent; FUJIFILM WAKO Chemicals, Japan) or on expanded vermiculite fertilised with 1/2,000 hyponex (HYPONEX Japan) every two weeks.

*A. agrestis* (Bonn strain) was grown on Half-strength Knop-II media (125 mg/L KNO3, 500 mg/L Ca(NO_3_)_2_ 4H_2_O, 125 mg/L Mg(SO_4_)_2_ 7H_2_O, 125 mg/L KH_2_PO_4_, and 10 mg/L FeCl_3_ 6H_2_O; pH5.5) containing 1% agar (FUJIFILM Wako Pure Chemical Corporation, Osaka, Japan).

For phosphate deficient treatment (Figs. 5a and 7g), Gemmae of *M. paleacea* or thallus of *A. agrestis* were pre-incubated on half-strength Gamborg’s B5 medium with 1.0 % agar for 2 weeks at 22 °C under continuous light. After 2 weeks, the thallus was transferred to the medium with or without phosphate and incubated for another week. Resultant plants were harvested and used for expression analysis.

For low phosphate treatment (Figs. 5e, 6e-f and 7h), Gemmae of *M. paleacea* were incubated on half-strength Gamborg’s B5 medium with 1.0 % agar with reduced phosphorus concentration (5-fold or 10-fold low phosphate concentration) for the appropriate time as described in each legend.

To analyze thallus phenotypes, gemmae were grown on half-strength Gamborg’s B5 medium with 1.0% agar or the medium with a reduced phosphorus concentration, at 22 °C under continuous light for 21 days. Resultant plants were used for the analysis of thallus phenotypes. To accurately measure the area and rhizoid length, plants were flattened with a coverslip on a glass slide, and images were taken. To measure the rhizoid length, the average lengths of the three longest rhizoids of each thallus were used as a representative rhizoid length for each thallus. To measure the angle between the thallus and the growth medium, images were taken from the side. For all measurements, the images were analyzed using ImageJ^44^.

### Plasmid construction and plant transformation

To construct vectors to generate CRISPR-Cas9 mutants, oligo DNAs containing the target sequences and the complementary sequences were annealed and introduced into pMpGE_En01 (*ccd8a, ccd8b, kai2a, kai2b*) or pMpGE_En03 (*max1, max2*). Subsequently, they were inserted into pMpGE010 (*ccd8b-1, -2, kai2a-2, kai2b-1, -2*) or pMpGE011 (*ccd8a-1, -3, max1-1, -2, kai2a-1, max2-1, -2*) by Gateway LR reactions. Transformation of *M. polymorpha* was performed as described^67^, and slight modifications were applied for the transformation of *M. paleacea*. For transformation of *M. polymorpha*, 10- to 14-day-old thalli grown from gemmae were transferred to fresh media and incubated for 2–3 days before *Agrobacterium* infection. For transformation of *M. paleacea*, 1- to 2-month-old thalli were cut and incubated on fresh media for 5–7 days before infection. To generate the Mpa*kai2kai2b* double mutant, *KAI2A* guide RNA inserted in pMpGE011 was introduced to *kai2b-1*. To generate the Mpa*ccd8accd8b* double mutant, *CCD8A* guide RNA inserted in pMpGE011 and *CCD8B* guide RNA inserted in pMpGE010 were introduced to WT thalli at the same time (Mpa*ccd8a-2/8b-3*) or *CCD8B* guide RNA inserted in pMpGE010 was introduced into *ccd8a-1* (Mpa*ccd8a-1/8b-4*).

### Analysis of CCD8A in *Marchantiopsida* species

Genomic DNA of *Marchantia chenopoda*, *M. emarginata* and *M. pinnata* was extracted as reported previously^68^. 2.5 kb of the CCD8A gene was PCR-amplified using GoTaq polymerase (Promega) according to the manufacturer’s protocol and the primers MpCCD8-F and MpCCD8-R (Supplementary Table 5). Purified PCR products were Sanger-sequenced at Eurofins (Germany) using the same primers.

### SL analysis in exudates and plants

*M. paleacea*, *M. polymorpha*, *M. pinnata*, *M. emarginata* and *P. carolinianus* were grown in 10 cm diameter pots filled with vermiculite under a 15 h light (50 μmol m^−2^ s^−1^)/9 h dark photoperiod at 23 °C in a growth room. *D. erythrosora* and *M. struthiopteris* plants were grown in a glasshouse with temperature control and under a natural photoperiod, at 23 °C in the day and 18 °C at night during spring to summer. *A. officinalis* and *A. hypogaea* seedlings were grown at 25 °C in the day and 20 °C at night. These plants were grown using 1/2,000 Hyponex and tap water. At least two weeks after growing in tap water, 200 ml per pot of water was poured on the plants (*Marchantia* and *Phaeoceros*) or the surface of the vermiculite (other plants), and collected from the bottom of the pot. The exudates were extracted with ethyl acetate. For SL quantification, 5 ng each of [1-^13^CH_3_]*rac*-carlactonoic acid and [3a,4,4,5,5,6′-D_6_]4DO were added as internal standards prior to the extraction. Endogenous SLs in *M. paleacea* plants were extracted with acetone. SLs were analyzed by LC-MS/MS using QTRAP 5500 (AB Sciex, USA) as reported previously^21^. *rac*-CL, *rac*-CLA, [1-^13^CH_3_]*rac* -CLA and [3a,4,4,5,5,6′-D_6_]4DO were synthesized as described previously^69^.

### Seed germination assay

*Orobanche minor* seeds were collected from mature plants that parasitized red clover (*Trifolium pretense* L.) grown in Tochigi Prefecture, Japan in June 2017. Germination assay of *O. minor* seeds was conducted as reported previously^50^. *Phelipanche ramosa* and *Striga hermonthica* seeds were kindly supplied by Profs. Philippe Delavault (University of Nantes, France) and A. G. T. Babiker (Sudan University of Science and Technology, Sudan), respectively. Seeds of *P. ramosa* and *S. hermonthica* were surface-sterilized in 1% NaOCl containing 0.1% Tween-20 for 5 min. The seeds were then rinsed 10 times with sterile distilled water and air-dried. Twenty to 40 seeds were sown on a 5 mm-glass fiber disc prepared by a hole puncher (Whatman GF/A, UK). Twenty discs were placed in a 5 cm-Petri dish lined with two layers of filter paper wetted with 1.5 ml of distilled water. The Petri dishes were sealed with parafilm, enclosed in polyethylene bags and incubated in the dark at 18°C for 7 days (*P. ramosa*) or at 30°C for 15 days (*S. hermonthica*). After conditioning, the glass fiber discs were blotted to remove excess water. Each group of three discs was transferred to a separate 5 cm-Petri dish lined with two layers of filter paper and wetted with 0.6 ml of test solutions. The Petri dishes were sealed, enclosed in polyethylene bags, and placed in the dark at 20°C for 6 days (*P. ramosa*) or at 30°C for 3 days (*S. hermonthica*). The germination rate was determined under a 20× binocular dissecting microscope.

### Isolation and purification of bryosymbiol (BSB)

*M. paleacea* was grown on vermiculite in a plastic container (with holes in the bottom, 38×25×14 cm, W×L×H) using tap water. Eighteen containers were placed under a 14 h light (60 μmol m^−2^ s^−^1)/10 h dark photoperiod at 24 °C in a growth room. Water was poured on the plants and circulated every 10 min per hour using an aquarium pump. The exudates released into the water were adsorbed on the activated charcoal (2 g per 3 L) in the pump. Fresh activated charcoal was replaced every 2 d. The collected charcoal was eluted with acetone. After evaporation of the acetone *in vacuo*, the aqueous residue was extracted with EtOAc. The EtOAc phase was washed with 0.2 M K_2_HPO_4_ (pH 8.3), dried over anhydrous MgSO_4_, and concentrated *in vacuo*. The concentrated samples were kept at 4 °C until use. The crude EtOAc extract (107.1 mg), collected over five weeks, was subjected to silica gel column chromatography (55 g) with stepwise elution of *n*-hexane-EtOAc (100:0 to 0:100, v/v, 10 % step) to give fractions 1–11. Fractions 6 (50% EtOAc, 13.2 mg) containing a novel SL was subjected to silica gel column chromatography (20 g) using *n*-hexane-EtOAc (60:40, v/v). Fractions were collected every 10 ml. Fractions 9–18 were found to contain a novel SL based on LC-MS/MS and GC-MS analyses. Fractions 9–18 were combined (3.17 mg) and purified by HPLC on an ODS column (Kinetex C18, 4.6×250 mm, 5 μm; Phenomenex, USA) with a MeCN/H_2_O gradient system (30:70 to 100:00 over 40 min) at a flow rate of 1 ml/min. The column temperature was set to 30 °C. The fraction eluted as a single peak at 28.6 min (detection at 248 nm) was collected. This fraction was further purified by HPLC on a Develosil ODS-CN column (4.6×250 mm, 5 μm; Nomura Chemicals, Japan) with isocratic 70% MeCN/H_2_O at a flow rate of 0.8 ml/min to afford BSB (0.33 mg, *Rt* 21.1 min, detection at 248 nm).

### Structural determination of BSB

HR–ESI–TOF–MS (AB Sciex TripleTOF 5600) showed a protonated molecular ion at *m/z* 349.1655 [M+H]^+^ calcd for C_19_H_25_O_6_, *m/z*: 349.1651). EI-GC/MS (JEOL JMS-Q1000GC/K9 on an Agilent DB-5 column) gave a molecular ion peak at *m/z* 348 [M]^+^, with fragment ion peaks at *m/z* 235, 217, 165 (the base peak), 123, and 97 (the D-ring fragment).

The ^1^H and ^13^C NMR spectra (JEOL JMN-ECA-500 spectrometer in CDCl_3_ (δ_H_ 7.26, δ_C_ 77.0)) aided with DEPT and HMQC experiments (Supplementary Table S3) revealed 19 carbon signals including two α,β-unsaturated ester carbonyls, two trisubstituted olefins, one tetrasubstituted olefin, three sp^3^ oxymethines, three sp^3^ <u>methylenes</u>, two methyls, one quaternary sp^3^ carbon, and two allylic methyls.

The ^1^H-^1^H COSY and HMBC correlations from H-2′, H-3′, H-6′, and H-7′ (Supplementary Fig. 3a-c and Supplementary Table 1) indicated the presence of the conserved enol ether-bridged methylbutenolide (the D-ring). The HMBC correlations of H-6′ to C-2, C-3, and C-4 and of H-5 to C-4 established the presence of α-methylene-β-hydroxy-γ-butyrolactone ring (the C-ring). The ^1^H-^1^H COSY and HMBC correlations from three methyls at H-12, H-13, and H-14 and from three sp^3^ methylenes at H-8, H-9, and H-10 showed a 2,6,6-trimethylcyclohex-1-en-1-yl moiety (the A-ring). The two substructures, the A- and the CD-parts, were connected together at C-5 and C-6 as indicated by the HMBC correlation of H-5 to C-6 and C-11. Therefore, the chemical structure of BSB was determined as (*E*)-5-((4-hydroxy-2-oxo-5-(2,6,6-trimethylcyclohex-1-en-1-yl)dihydrofuran-3(2*H*)-ylidene)methoxy)-3-methylfuran-2(5*H*)-one.

The CD spectrum (JASCO J-820W spectropolarimeter in acetonitrile) had a positive and negative Cotton effect around 224 nm (Δε 27.4) and 266 nm (Δε −4.44), respectively (Supplementary Fig. 3d), indicating that BSB has a 2′(*R*) configuration as previously identified natural SLs. The NOESY correlations between H-12/H-13 methyls and H-5 oxymethine, and between H-14 methyl and H-4 oxymethine suggested the relative stereochemistry at C-4 and C-5 is *anti*. Accordingly, we tentatively assigned the stereochemistry to be (4*R*, 5*S*, 2′*R*) (Fig. 2f).

### Expression of Mpa*MAX1* in yeast

Total RNAs were extracted from whole *M. paleacea* plants with rhizoids using the RNeasy Plant Mini Kit (Qiagen, Germany) and used to synthesize single-stranded cDNAs with the SuperScript III First-Strand Synthesis System (Invitrogen, USA). PCR amplification was performed with PrimeSTAR HS DNA polymerase (TaKaRa Bio, Japan) using the primers MpaMAX1_cDNA_F and MpaMAX1_cDNA_R (Supplementary Table 5). The MpaMAX1 cDNA was cloned into the *BamH I* and *Kpn I* sites of the pYeDP60 vector with the GeneArt Seamless Cloning and Assembly Enzyme Mix (Invitrogen, USA) using the primers MpaMAX1_cloning_F and MpaMAX1_cloning_R (Supplementary Table 5). At least four clones were sequenced to check for errors in the PCR. The accession number of MpaMAX1 was deposited as LC553047. Heterologous expression of MpaMAX1 in yeast (*Saccharomyces cerevisiae*) was carried out as described previously^21, 69^. Microsomes (100 μl) were incubated with *rac*-CL (0.25 to 33 μM) and NADPH (500 μM) at 28 °C for 10 min. The reaction mixtures were extracted with ethyl acetate, dried over anhydrous Na2SO4 and then subjected to LC-MS/MS analysis. The product BSB was quantified using the peak area in the transitions of *m/z* 349 to 97 by assuming the same ion intensity as that of 4-deoxyorobanchol. The kinetics parameter was determined using triplicate samples and calculated by the Michaelis-Menten equation using SigmaPlot 14 (Systat Software, USA). The *V*_max_ of BSB from *rac*-carlactone was 92.8 ± 8.4 nM min^−1^ and the Michaelis constant was 4.78 ± 1.22 μM.

### Mycorrhization assays

Thalli of *Marchantia paleacea*, *Marchantia emarginata*, *Marchantia pinnata* and *Marchantia chenopoda* were grown on a substrate composed of 50 % zeolite and 50 % sand (0.7–1.3 mm) in 7×7×8 cm pots (five thalli per pot). Each pot was inoculated with *ca*. 1,000 sterile spores of *Rhizophagus irregularis* DAOM 197198 (Agronutrition, Labège, France) and grown with a 16 h/8 h photoperiod at 22 °C/20 °C. Pots were watered once per week with Long Ashton medium containing 7.5 μM of phosphate^30^. For strigolactone complementation assays, 10 ml of Long Ashton solution with 10^−7^ M GR24 or 0.1 % acetone (control) was added to each pot weekly, starting on the day of the inoculation. Hyphal branching activity of BSB on *Gigaspora margarita* was evaluated as reported previously^33^.

To establish the mycorrhizal status of the different Marchantia species, thalli were embedded in 6 % agarose and 0.1 mm sections were prepared using a Leica vt1000s vibratome. Sections were incubated overnight in PBS containing 1 μg/ml WGA-FITC (Sigma) and pictures were taken with a Zeiss Axiozoom V16 microscope.

For the quantitative evaluation of symbiosis, thalli were harvested six weeks post-inoculation (*Marchantia paleacea* mutants) or eight weeks post-inoculation (diverse Marchantia species). For the analysis of the mycorrhizal phenotype of the mutants, longitudinal hand sections were cleared in 10 % KOH overnight and ink coloured for 30 min in 5 % Sheaffer ink, 5 % acetic acid. Sections were observed under a microscope and infection points were scored for each thallus.

### Phylogenetic analysis

Homologues of the reference genes *CCD8* and *MAX1* of *Arabidopsis thaliana* or *Medicago truncatula* were retrieved in a custom database composed of 197 species covering the main land plant lineages (Supplementary Table 1) and including, for Lycophytes, Monilophytes and Bryophytes, predicted coding sequences of transcriptomic data from the 1KP project^16, 57^. Searches were perform using the tblastn v2.9.0+ algorithm with default parameters and an e-value threshold of 1^e-10^.

For each gene, their putative homologue coding sequences were aligned using MAFFT v7.313 with default parameters and the resulting alignments trimmed to remove positions with more than 80 % gaps using trimAl v1.4.1. Then, alignments were subjected to Maximum Likelihood (ML) analysis using IQ-TREE v1.6.7 and branch support tested with 10,000 replicates of SH-aLRT and UltraFast Bootstraps. Prior to ML analysis, the best-fitting evolutionary model was tested using ModelFinder and retained according to the Bayesian Information Criteria. Trees were visualized and annotated using the iTOL v5.6.3^70^.

### RNA extraction and expression analysis

RNA was extracted from frozen plant samples using the NucleoSpin RNA Plant kit (MACHEREY-NAGEL). cDNAs were synthesized using SuperScript VILO (Invitrogen). Quantitative PCR was performed using the KOD SYBR®qPCR Mix (Toyobo Life Science) with Light Cycler 480 II (Roche). Primers used to amplify the cDNAs are shown in Supplemental Table 5. The ACTIN gene of each species was used as a standard.

### RNA-seq analysis

Gemmae of WT, Mpa*ccb8a-1/b-4*, and Mpa*ccb8a-2/b-3* were incubated on half-strength Gamborg’s B5 medium for 2 weeks at 22 °C under continuous light. After 2 week-culture, the thalli were transferred to the medium with or without phosphate. Three biological replicates were prepared for each genotype. Total RNA was extracted using the NucleoSpin RNA Plant kit (Macherey-Nagel, Germany). mRNA was isolated from the total RNA with an NEB Next poly(A) mRNA Magnetic Isolation Module Kit (New England Biolabs, MA, USA). RNA-seq libraries were synthesized using an NEB Next Ultra RNA Library Prep Kit for Illumina and an NEB Next Adaptor for Illumina Kit (New England Biolabs, MA, USA). Paired-end sequencing was performed using the HiSeqXten (Illumina, CA, USA).

The reads were mapped onto the *Marchantia paleacea* genome^48^ assembly using HISAT2^71^ for Galaxy (https://usegalaxy.org) with default parameters. Read counts were calculated by Feature Counts for Galaxy. To identify differentially expressed genes, q-values were calculated by the TCC R package^72^, and genes with q-value < 0.05 and log2-fold change > 1 or log2-fold change < −1 were selected as up-or down-regulated genes. Summary statistics of RNA-seq analysis are available in Supplementary Table 4.

### Differential Scanning Fluorimetry (DSF) analysis

The coding sequences for MpaKAI2A and MpaKAI2B were amplified from cDNA synthesized from total mRNA of the *Marchantia paleacea* thalli using the following primer sets, MpaKAI2A_cDNA_F and MpaKAI2A_cDNA_R for MpaKAI2A and MpaKAI2B_cDNA_F and MpaKAI2B_cDNA_R for MpaKAI2B (Supplementary Table 5). The PCR products were cloned into the modified pET28 vector using the In-Fusion HD Cloning Kit (Clontech) to generate pET28M-MpaKAI2A and pET28M-MpaKAI2B. *Escherichia coli* strain BL21 (DE3) (Takara) was used for recombinant protein expression. Overnight cultures (3 ml) in LB liquid medium containing 100 μg/ml kanamycin were inoculated into fresh LB medium (1 L) containing 100 μg/ml kanamycin and incubated at 37 °C until the OD600 value reached to 0.6 ~ 0.8. Then, IPTG was added to 200 μM, and the cells were further incubated at 25 °C overnight. The cells were collected by centrifugation at 4000 rpm for 15 min. The pellets were resuspended in 30 ml of buffer A (20 mM Tris-HCl (pH 8.0) and 200 mM NaCl). The cells were crushed using a cell disruption device (UD-200, TOMY SEIKO). The soluble fraction was loaded on Ni Sepharose 6B fast flow (Cytiva) pre-equilibrated with buffer A. The bound MpaKAI2A and MpaKAI2B proteins were eluted with 5 ml of buffer A containing 125 mM of imidazole. The purified proteins were concentrated using an Amicon Ultra-4 10 K (Millipore).

For DSF analysis, 20 μl reaction mixtures containing 10 μg protein, 0.015 μl Sypro Orange and GR24s with final concentrations of 250 μM, 100 μM or 50 μM, were prepared in 96-well plates. The final acetone concentration in the reaction mixture was 5%. 1×PBS buffer (pH 7.4) containing 5% acetone was used in the control reaction. DSF experiments were performed using a Light Cycler 480 II (Roche). Sypro Orange (Invitrogen) was used as the reporter dye. Samples were incubated at 25 °C for 10 min, then, the fluorescence wavelength was detected continuously from 30 °C to 95 °C. The denaturation curve was obtained using MxPro software (Agilent).

### Generation of AtD14-overexpressing plants

To generate *AtD14*ox lines, the AtD14 coding sequence cloned into pENTR/D-TOPO vector^40^ was inserted into pMpGWB103 or pMpGWB303 by Gateway LR reactions. The MpEFpro: AtD14 vector was introduced into regenerating thalli of Tak-1 or WT of *M. paleacea* as described above.

To generate Mp*AtD14*ox/Mp*kai2a* or Mpa*AtD14*ox/Mpa*kai2a* line, *MpKAI2A* or *MpaKAI2A* was knocked out in Mp*AtD14*ox or Mpa*AtD14*ox lines. *MpKAI2A* guide RNA inserted in pMpGE010^44^ or *MpaKAI2A* guide RNA inserted in pMpGE011 were introduced into Mp*AtD14*ox or Mpa*AtD14*ox lines, respectively.

### Application of GR24

Synthetic strigolactone analog *rac*-GR24 was obtained from Chiralix B.V. (Netherland). The enantiomers of GR24 were obtained by optical resolution from *rac*-GR24 stationary phase (Chiralpak IC-3, 250 × 4.6 mm, Daicel, Japan) under normal-phase using hexane/isopropanol (70/30) by HPLC. GR24^4DO^ and GR24^5DS^ were identified by CD analysis. *rac*-GR24, GR24^4DO^, and GR24^5DS^ dissolved in acetone were added to half-strength Gamborg’s B5 medium with 1.0 % agar. Thalli grown from gemmae for 2 weeks were incubated on the *rac*-GR24, GR24^4DO^, GR24^5DS^, or mock containing media for 6 hours.

### Statistical analysis

We used Welch’s t test or Tukey’s honestly significant difference (HSD) test to evaluate statistical significance. Experimental sample sizes and statistical methods detail are given in the each legend.

## Data availability

All reads obtained have been deposited in the DDBJ and are available through the Sequence Read Archive (SRA) under the accession numbers DRA012476. The authors declare that all data supporting the findings of this study are available within the manuscript and its supplementary files or are from the corresponding author upon request.

## Acknowledgements

We thank Philippe Delavault (University of Nantes) and A.G.T. Babiker (Sudan University of Science and Technology) for *Phelipanche ramosa* and *Striga hermonthica* seeds, respectively. We are grateful to the genotoul bioinformatics platform Toulouse Occitanie (Bioinfo Genotoul, doi: 10.15454/1.5572369328961167E12) for providing computing resources. Researches conducted by Ju.K. lab and Ta.N. lab were supported by a Grants-in-Aid from the Ministry of Education, Culture, Sports, Science, and Technology, Japan (20H05684 and 17H06475 to Ju.K., 19K05838 to Ta.N., 18K05452 to X.X.) and Canon Foundation to Ju.k. Research conducted in P.-M.D. lab was supported by the Agence Nationale de la Recherche (ANR) grant EVOLSYM (ANR-17-CE20-0006-01) and was supported by the project Engineering Nitrogen Symbiosis for Africa (ENSA) currently supported through a grant to the University of Cambridge by the Bill & Melinda Gates Foundation (OPP1172165) and the UK Government’s Department for International Development (DFID). The Laboratoire de Recherche en Sciences Végétales (LRSV) belongs to the TULIP Laboratoire d’Excellence (ANR-10-LABX-41).

## Author contributions

Ju.K., P.-M.D., M.K.R. and Ta.N. designed the research and wrote the article. K.K., M.K.R., S.S., Y.M. and A.K. produced mutants and analyzed phenotypes. K.K., S.S., Y.M., A.K., Y.L., H.S. and H.K. examined gene expression. S.S., M.K.R., To.Na. and P.-M.D. analyzed A.M. symbiosis. C.L. and Je.K. conducted phylogenetic analysis. S.S., A.Y., K.Y., K.M., S.Y., Y.K. and Y.T. analyzed BSB and enzyme activity. X.X., K.U. and K.A. determined the chemical structure of BSB. K.S., M.S., To.Na. and To.Ni. prepared materials.

## Additional information

### Supplementary information

Supplementary Figure 1: Phylogenetic analysis of *CCD8* genes.

Supplementary Figure 2: Phylogenetic analysis of *MAX1* genes.

Supplementary Figure 3: 1D and 2D NMR spectra (CDCl3), and CD spectra of bryosymbiol.

Supplementary Figure 4: Strigolactones in hornwort, moss, ferns and seed plants.

Supplementary Figure 5: Mycorrhizal phenotype of strigolactone biosynthesis and signalling mutants.

Supplementary Figure 6: Evaluation of the MpaKAI2-SLs interaction.

Supplementary Figure 7: Effects of SLs and phosphorus on gene expressions in *AtD14* line in *M. polymorpha* and in *M. paleacea*.

Supplementary Table 1: Species used in the analysis shown in Supplementary Figs 1 and 2.

Supplementary Table 2: Production of CRISPR mutants.

Supplementary Table 3: NMR spectroscopic data for BSB.

Supplementary Table 4 (Excel file): Results of RNAseq analysis.

Supplementary Table 5: Oligonucleotide primers used in this study.

**Correspondence** and requests for materials should be addressed to Ju.K., Ta.N., or P.-M.D.

