## Supplemental Figs and Tables for "An Ancestral Function of Strigolactones as Symbiotic Rhizosphere Signals"

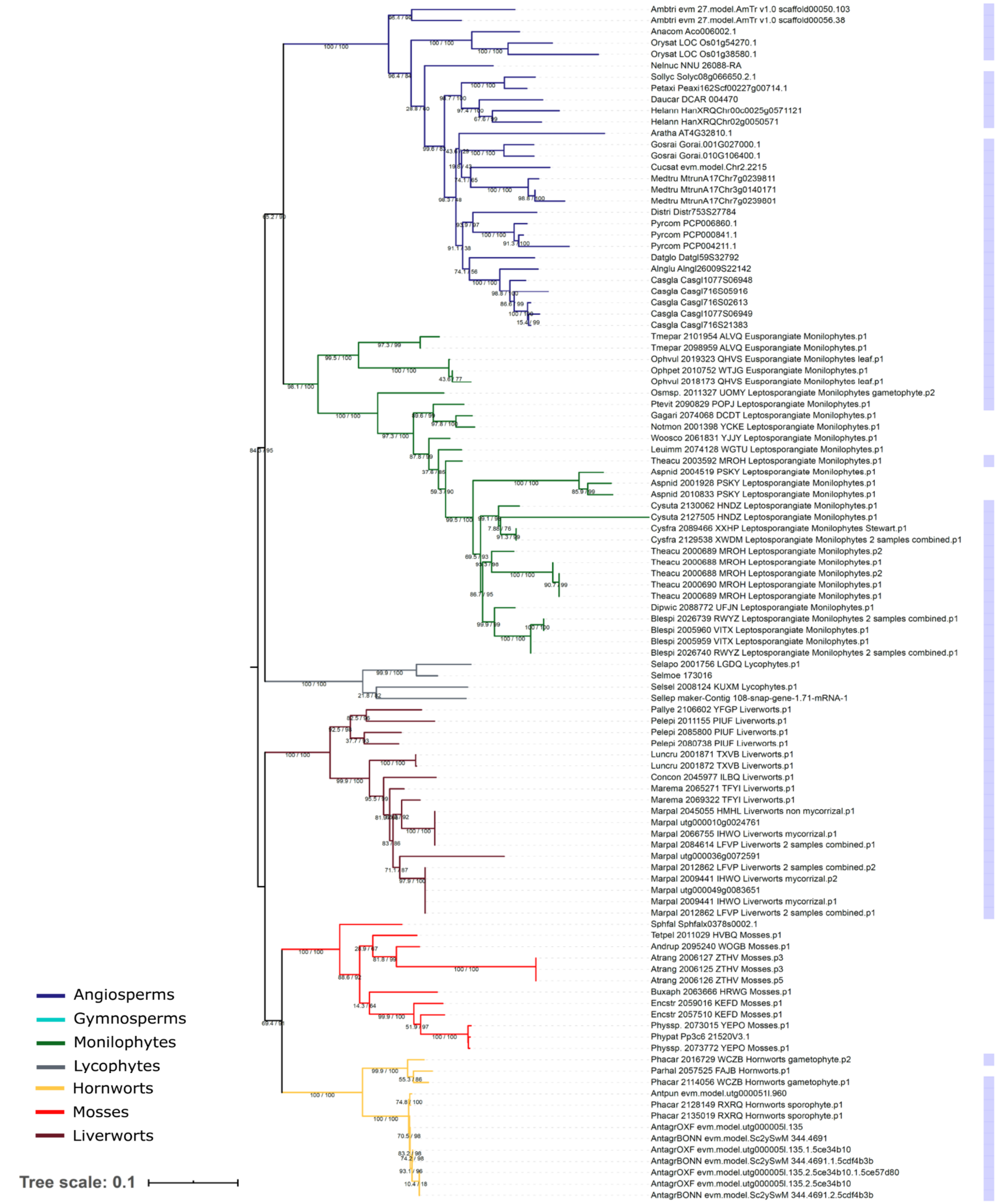


**Supplementary Figure 1. Phylogenetic analysis of *CCD8* genes.** Tree CCD8: Maximum Likelihood of *CCD8* gene (model: SYM+R5; log-likelihood: -57366.2677). The tree is rooted on the divergence node between vascular and non-vascular plants. Cyan boxes at the right of the tree mark species able to form arbuscular mycorrhizal symbiosis. Branches are coloured according to plant lineages. SH-aLRT and UltraFast Bootstraps branch supports are indicated by the number below the branches on both sides of the “/” symbol.

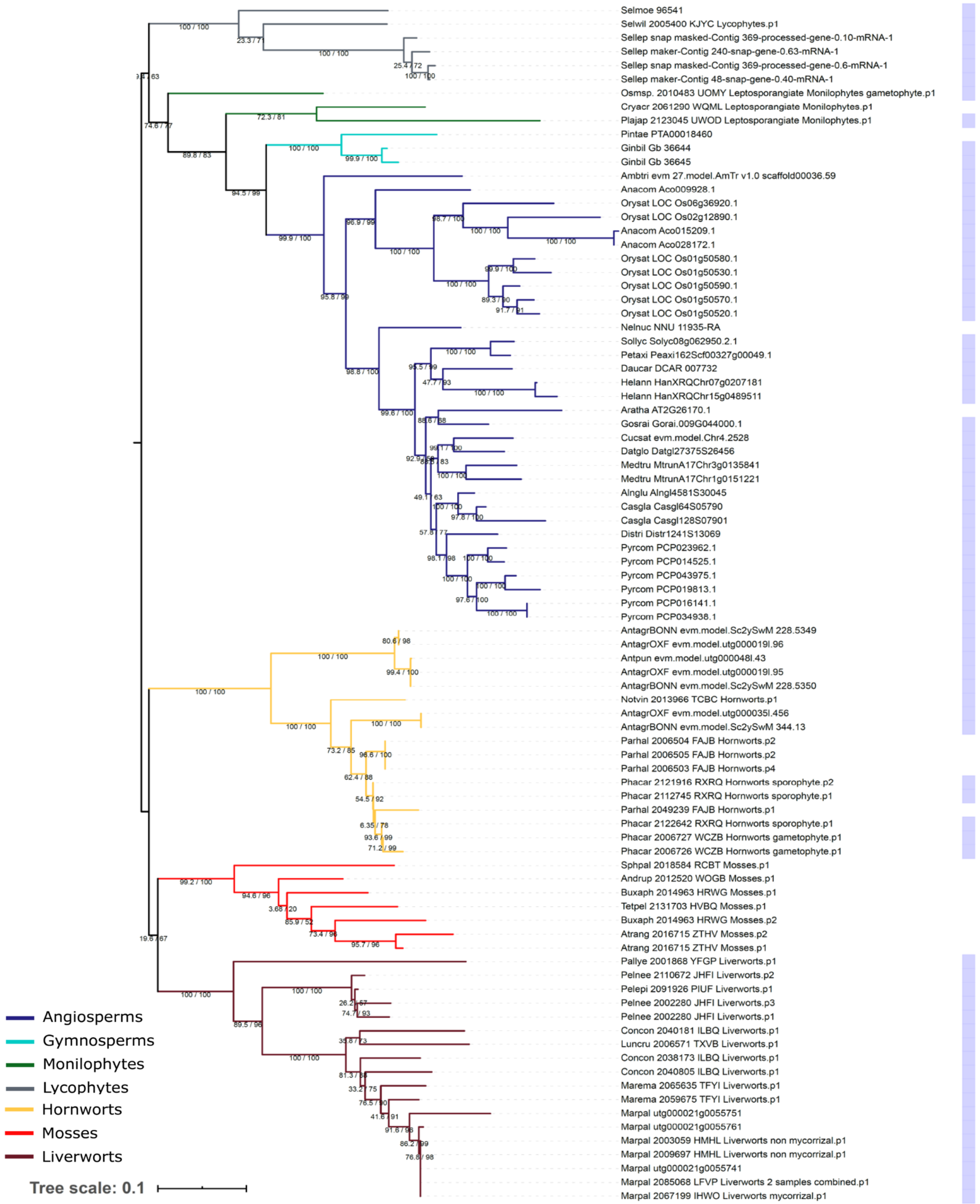


**Supplementary Figure 2. Phylogenetic analysis of *MAX1* genes.** Maximum Likelihood of *MAX1* gene (model: TVMe+R4; log-likelihood: -55502.0476). The tree is rooted on the divergence node between vascular and non-vascular plants. Cyan boxes at the right of the tree mark species able to form arbuscular mycorrhizal symbiosis. Branches are coloured according to plant. SH-aLRT and UltraFast Bootstraps branch supports are indicated by the number below the branches on both sides of the “/” symbol.

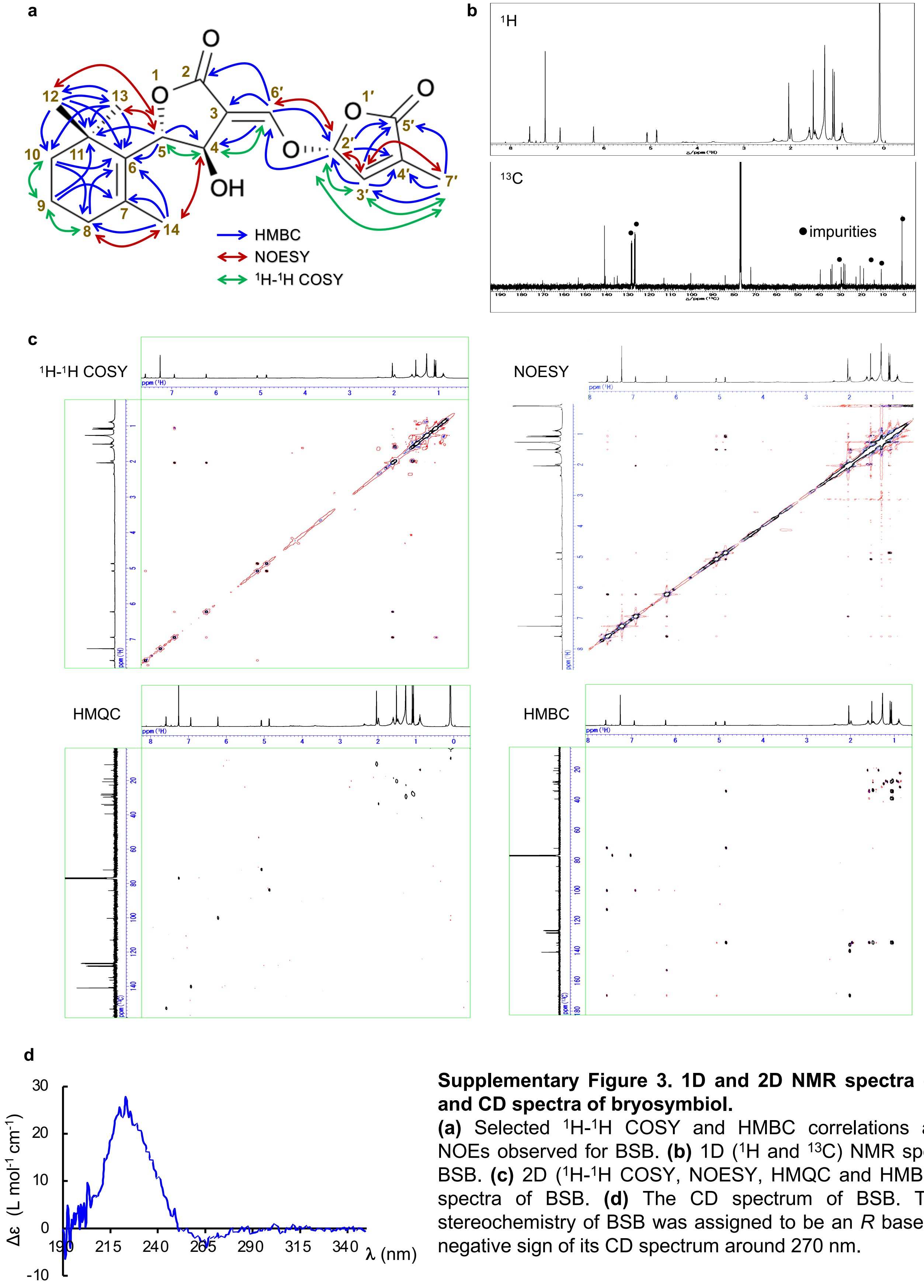


**Supplementary Figure 3. 1D and 2D NMR spectra ($CDCl_3$), and CD spectra of bryosymbiol.**
**(a)** Selected $^1$H-$^1$H COSY and HMBC correlations and key NOEs observed for BSB. **(b)** 1D ($^1$H and $^{13}$C) NMR spectra of BSB. **(c)** 2D ($^1$H-$^1$H COSY, NOESY, HMQC and HMBC) NMR spectra of BSB. **(d)** The CD spectrum of BSB. The C-2' stereochemistry of BSB was assigned to be an *R* based on the negative sign of its CD spectrum around 270 nm.

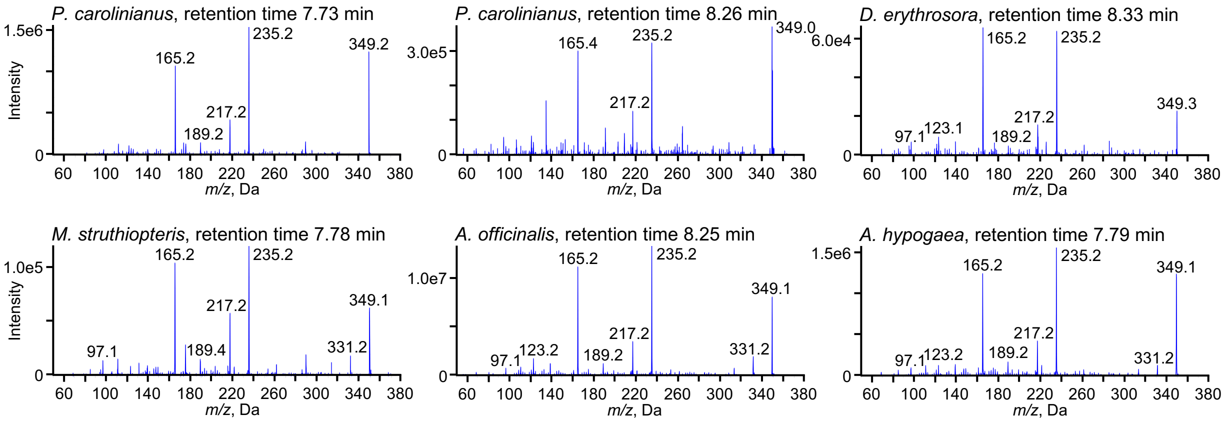


**Supplementary Figure 4. Strigolactones in hornwort, moss, ferns and seed plants.**
Product ion spectra of BSB. Product ion spectra derived from the precursor ion (*m/z* 349 in positive mode) of peaks detected in hornwort, ferns and seed plants are shown.

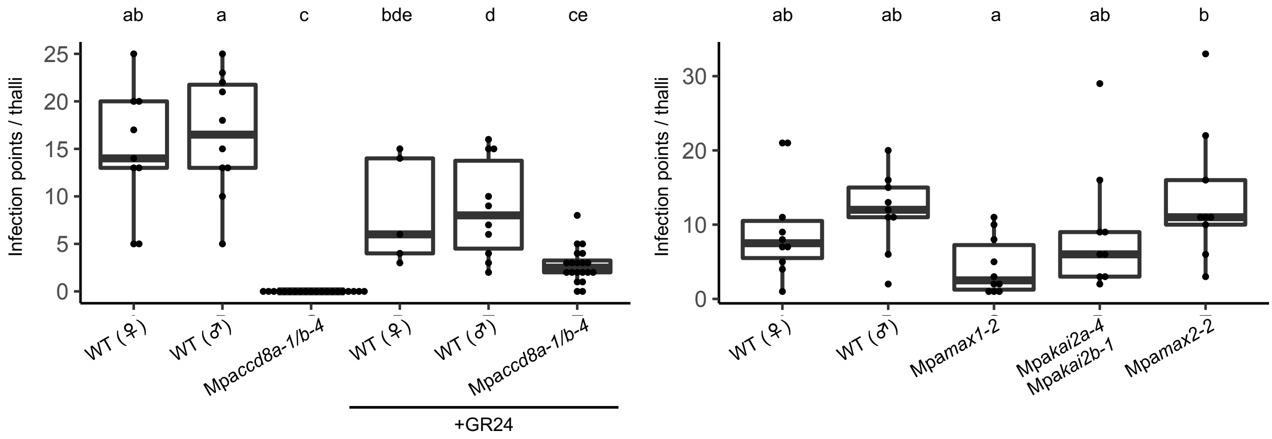


**Supplementary Figure 5. Mycorrhizal phenotype of strigolactone biosynthesis and signalling mutants.** Replication of experiments shown in Figure 5c conducted in independent mutant alleles is shown (n ≥ 5). The number of infection points per thalli in BSB biosynthesis and signalling mutants of *M. paleacea*. Letters show different statistical groups (ANOVA, post hoc Tukey).

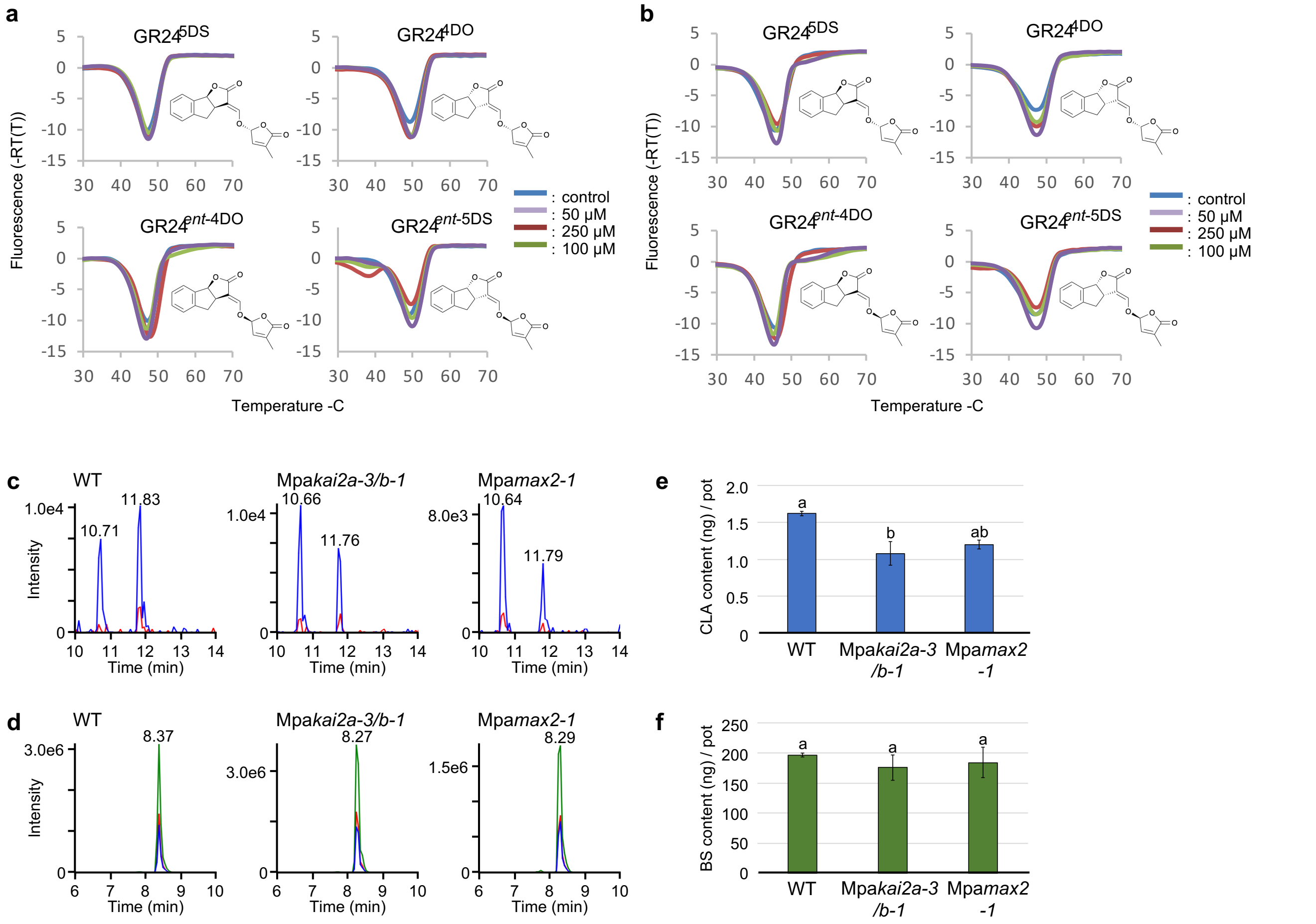


**Supplementary Figure 6. Evaluation of the MpaKAI2-SLs interaction.**

**(a and b)** Differential Scanning Fluorimetry (DSF) analysis with 4 stereoisomers of GR24 and MpaKAI2A (a) and MpaKAI2B (b) protein. **(c and d)** Detection of carlactonoic acid and BSB in exudates of the Mp*akai2a/b* and Mp*amax2* mutants. MRM chromatograms of carlactonoic acid (c) and BSB (d) are shown. **(e and f)** The content of carlactonoic acid and BSB in exudates of the Mp*akai2a/b* and Mp*amax2* mutants. The exudates of WT, Mp*akai2a/b* and Mp*amax2* grown in 10 cm diameter pots were analysed by LC-MS/MS. Carlactonoic acid (e) was quantified using [1-$^{13}CH_3$] carlactonoic acid as an internal standard. BSB (f) was quantified using [$^{2}H_6$]4DO as an internal standard. Data are the means ± SD (n = 3, ANOVA with Tukey's test, $p < 0.05$).

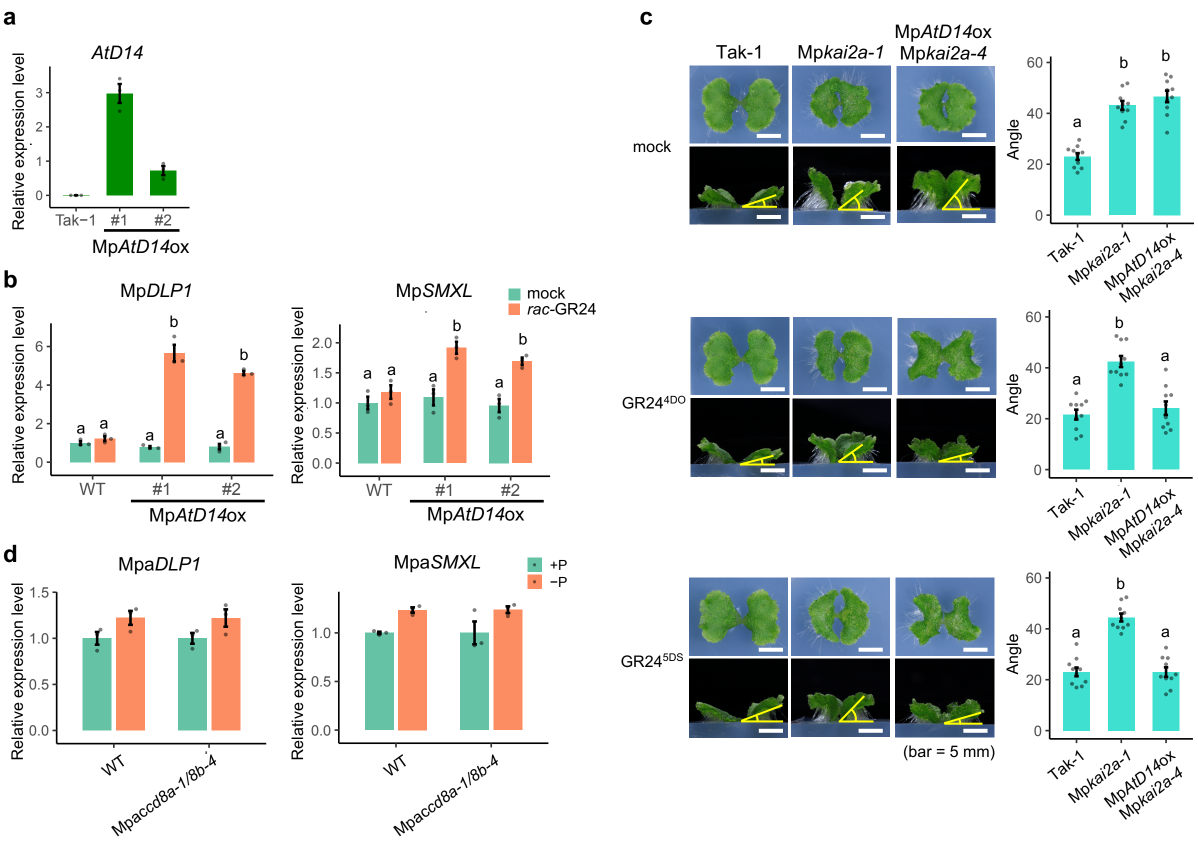


**Supplementary Figure 7. Effects of SLs and phosphorus on gene expressions in *AtD14* line in *M. polymorpha* and in *M. paleacea*.**

**(a)** Expression of *AtD14* introduced into *M. polymorpha*. **(b)** Expression of Mp*DLP1* and Mp*SMXL*, markers of the KAI2-dependent signaling pathway, in WT and Mp*AtD14*ox lines of *M. polymorpha* after *rac*-GR24 treatment. **(c)** Complementation of Mp*kai2a* phenotypes in Mp*AtD14*ox/Mp*kai2a* (*M. polymorpha*) line by the addition of 1 µM GR24$^{4DO}$ or 1 µM GR24$^{5DS}$ **(d)** Expression of Mpa*DLP1* and Mpa*SMXL* after phosphate starvation in WT and Mpa*ccd8a-1/8b-4*.

Values in (a), (b) and (d) are means ± SD (n = 3), and values in (c) are means ± SD (n = 10). The HSD test was used for multiple comparisons and statistical differences (P-values < 0.05) are indicated by different letters.

### Supplementary Table 2. Production of CRISPR mutants

| Allele | gRNA sequence | Mutation site | Amino acid |
|---|---|---|---|
| Mpa*kai2a-1* | GGACTGACCAGTCTGTGTGGA | c.93_94insT | p.Trp32LeufsTer11 |
| Mpa*kai2a-2* | TGTACGATACCATGGGAGCA | c.157_158delinsA | p.Ala53LysfsTer28 |
| Mpa*kai2a-3* | TGTACGATACCATGGGAGCA | c.156_157del | p.Ala53ArgfsTer22 |
| Mpa*kai2a-4* | TGTACGATACCATGGGAGCA | c.156del | p.Ala53GlnfsTer28 |
| Mpa*kai2a-5* | GGACTGACCAGTCTGTGTGGA | c.93_94insT | p.Trp32LeufsTer11 |
| Mp*kai2a-4* | GGAGATCAGCTCGTGGTACT | c.55_59del | p.Val19ThrfsTer8 |
| Mpa*kai2b-1* | TTGGCACGGATCAGTCAGTG | c.88_91delinsAT | p.Ser30IlefsTer19 |
| Mpa*kai2b-2* | TTGGCACGGATCAGTCAGTG | c.1-104_117delins CAGGTGAGGTATAAGAT GCTTCCAAACA | No start codon |
| Mpa*ccd8a-1* | GGAACAGTGGGAAGGAGAGC | c.389_426+123del | p.Leu132GlufsTer16 |
| Mpa*ccd8a-2* | GTTCACAATAGCAGATAAAG | c.53_56del | p.Asp158LysfsTer16 |
| Mpa*ccd8b-1* | ACAATGGGAAGGAGAGCTGG | c.368_380delins AAATTCCACCGAGGAAT TGAGCCCTGGAC | p.Leu123GlnfsTer7 |
| Mpa*ccd8b-2* | ACAATGGGAAGGAGAGCTGG | c.367_368insT | p.Glu124GlyfsTer27 |
| Mpa*ccd8b-3* | ACAATGGGAAGGAGAGCTGG | c.365_369delins GAACTCAGGAATT | p.Glu122GlyfsTer19 |
| Mpa*ccd8b-4* | ACAATGGGAAGGAGAGCTGG | c.366_367insC | p.Leu123ProfsTer28 |
| Mpa*max1-1* | CTCTGCTAACCAAGCACC | c.157_63del | p.Lys54ArgfsTer19 |
| Mpa*max1-2* | ATGCAGAGCTGTGCCGAC | c.780_806del | p.Asp85ValfsTer428 |
| Mpa*max2-1* | ACCCAAATCAGCGACCTGC | c.19_58del | p.Aps11_Ile20delinsVal |
| Mpa*max2-2* | ACCCAAATCAGCGACCTGC | c.34_35del | p.Pro13ArafsTer19 |

※ Notations were followed den Dunnen JT and Antonarakis SE (2000). Hum.mutat. 15:7-12

**Supplementary Table 3. NMR spectroscopic data for BSB.**

| No. | δ $^{13}$C | δ $^{1}$H (mult., *J* Hz) | HMQC & DEPT | $^{1}$H–$^{1}$H COSY | HMBC | NOESY |
|---|---|---|---|---|---|---|
| 2 | 170.05 | | C | | | |
| *3* | 113.01 | | C | | | |
| *4* | 72.16 | 5.08 (dd, 4.6, 1.7) | CH | H-5, H-6′ | | H-14 |
| 5 | 84.20 | 4.87 (d, 4.6) | CH | H-4 | C-4, C-6 | H-12, H-13 |
| 6 | 134.88 | | C | | | |
| 7 | 134.97 | | C | | | |
| *8* | 33.95 | 1.986 (t, 6.1), 1.991 (t, 6.0) | $CH_2$ | H-9 | C-6 | H-14 |
| 9 | 19.10 | 1.57-1.63 (m) | $CH_2$ | H-8, H-10 | C-7, C-11 | |
| 10 | 39.48 | 1.44-1.51 (m) | $CH_2$ | H-9 | C-6, C-8 | |
| 11 | 34.55 | | C | | | |
| 12 | 28.50 | 1.06 (s) | $CH_3$ | | C-6, C-10, C-11, C-13 | H-5 |
| 13 | 27.86 | 1.10 (s) | $CH_3$ | | C-6, C-10, C-11, C-12 | H-5 |
| 14 | 20.75 | 1.51 (s) | $CH_3$ | | C-6, C-7, C-8 | H-4, H-8 |
| 2’ | 100.37 | 6.22 (quin, 1.4) | CH | H-3′, H-7′ | C-4′, C-5′, C-6′ | H-3′, H-6′ |
| 3’ | 140.57 | 6.94 (quin, 1.7) | CH | H-2′, H-7′ | C-2′, C-4′, C-5′ | H-2′, H-7′ |
| 4’ | 136.39 | | C | | | |
| 5’ | 169.88 | | C | | | |
| 6’ | 153.25 | 7.59 (d, 1.7) | CH | H-4 | C-2, C-3, C-4, C-2′ | H-2′ |
| *7’* | 10.79 | 2.04 (br.t, 1.7) | $CH_3$ | H-2′, H-3′ | C-3′, C-4′, C-5′ | H-3′ |

### Supplementary Table 5. Oligonucleotide primers used in this study.

| Use | Name | Sequence (5'→3') |
|---|---|---|
| Mutagenesis by CRISPR | MpaKAI2ACR1F | GCACCCAGCCTCTCGGACTGACCAGTCTGTGTGGAGTTTTAGAGCTAGAA |
| | MpaKAI2ACR1R | TTCTAGCTCTAAAACTCCACACAGACTGGTCAGTCCGAGAGGCTGGGTGC |
| | MpaKAI2ACR2F | GCACCCAGCCTCTCGTGTACGATACCATGGGAGCAGTTTTAGAGCTAGAA |
| | MpaKAI2ACR2R | TTCTAGCTCTAAAACTGCTCCCATGGTATCGTACACGAGAGGCTGGGTGC |
| | MpaKAI2BCR1F | GCACCCAGCCTCTCGTTGGCACGGATCAGTCAGTGGTTTTAGAGCTAGAA |
| | MpaKAI2BCR1R | TTCTAGCTCTAAAACCACTGACTGATCCGTGCCAACGAGAGGCTGGGTGC |
| | MpaCCD8ACR1F | GCACCCAGCCTCTCGGGAACAGTGGGAAGGAGAGCGTTTTAGAGCTAGAA |
| | MpaCCD8ACR1R | TTCTAGCTCTAAAACGCTCTCCTTCCCACTGTTCCCGAGAGGCTGGGTGC |
| | MpaCCD8ACR2F | GCACCCAGCCTCTCGGTTCACAATAGCAGATAAAGGTTTTAGAGCTAGAA |
| | MpaCCD8ACR2R | TTCTAGCTCTAAAACCTTTATCTGCTATTGTGAACCGAGAGGCTGGGTGC |
| | MpaCCD8BCR1F | GCACCCAGCCTCTCGACAATGGGAAGGAGAGCTGGGTTTTAGAGCTAGAA |
| | MpaCCD8BCR1R | TTCTAGCTCTAAAACCCAGCTCTCCTTCCCATTGTCGAGAGGCTGGGTGC |
| | MpaMAX1CR1F | CTCGCTCTGCTAACCAAGCACC |
| | MpaMAX1CR1R | AAACGGTGCTTGGTTAGCAGAG |
| | MpaMAX1CR2F | CTCGATGCAGAGCTGTGCCGAC |
| | MpaMAX1CR2R | AAACGTCGGCACAGCTCTGCAT |
| | MpaMAX2CR1F | CTCGACCCAAATCAGCGACCTGC |
| | MpaMAX2CR1R | AAACGCAGGTCGCTGATTTGGGT |
| Genotyping | MpaKAI2AgeF | CAATGTGCGGATAGTTGGTTCG |
| | MpaKAI2AgeR | TCTCAATAGATGCAAGGCACCC |
| | MpaKAI2BgeF | ACGTGTAGTAGGATCAGGGG |
| | MpaKAI2BgeR | GTCCGGTCTTTCCAAGGATG |
| | MpaCCD8AgeF | AGGAGAAAGCTGTAGAGATCGA |
| | MpaCCD8AgeR | CTCAGTTGGTTGTGAGATGCTA |
| | MpaCCD8BgeF | CACCCAAGTCCAGTGATTCG |
| | MpaCCD8BgeR | TCCCTCAAACTTCAAACGCA |
| | MpaMAX1geF | ATGGGGAGAGTTCGGCAGA |
| | MpaMAX1geR | CTGTCGTCTTAAAGCCCTCTG |
| | MpaMAX2geF | ATGGGGAGAGTTCGGCAGA |
| | MpaMAX2geR | CTGTCGTCTTAAAGCCCTCTG |
| qPCR | MpACTINqPCRF | AGGCATCTGGTATCCACGAG |
| | MpACTINqPCRR | ACATGGTCGTTCCTCCAGAC |
| | MpDLP1qPCRF | GGTGTGAAGAAAGTTGGAGTTATGG |
| | MpDLP1qPCRR | GTGTGAGGAATGAGGGATGGTT |
| | MpSMXLqPCRF | TGGGATGTCAGGTCGGAAAC |
| | MpSMXLqPCRR | AAAAGTTCTGCTGAGCTGCG |
| | MpaACTINqPCRF | ACGTGGCAATTCAGGCTGTC |
| | MpaACTINqPCRR | GCGTGGGGAAGAGCATAACC |
| | MpaDLP1qPCRF | ACCATCCGTCATTCCTCACAC |
| | MpaDLP1qPCRR | ATTTGTCTCCAATCTCAGGGTTCTT |
| | AtD14qPCRF | GCTTCGGTGGCGGAGTATCT |
| | AtD14qPCRR | AGGATGTTTCTGTGCCGGCT |
| | MpaD27qPCRF | GTACAAGGACTCTTGGCTTGAGAAA |
| | MpaD27qPCRR | GGAGCAAGTCGGAATGTGAAG |
| | MpaCCD7qPCRF | GAACCTGGAGAAGCAGGCTT |
| | MpaCCD7qPCRR | GGGGACTCTCAGTCTAGCCA |
| | MpaCCD8AqPCRF | CTATGGCTGGCTGGGACATATTG |
| | MpaCCD8AqPCRR | CTCTTCGCAGCATTGTACGC |
| | MpaCCD8BqPCRF | CTGCACTTGACCTTCCCGATG |
| | MpaCCD8BqPCRR | GTGGGAGGACAATCTTGGGG |
| | MpaMAX1qPCRF | ACATCGAGCTGGGTGGCTAC |
| | MpaMAX1qPCRR | TCGCTCGGGTCGAAATTCCT |
| | MpaSMXLqPCRF | TGCCACCATCAAGCACACCT |
| | MpaSMXLqPCRR | GAACAACGGGCTTTGCGTCA |
| | AaACTINqPCRF | GGCATCACACTTTCTACAATGAGC |
| | AaACTINqPCRR | TGACACCATCACCAGAATCAAGC |
| | AaCCD8qPCRF | TGTCAAGCAGCCGAGATGG |
| | AaCCD8qPCRR | GTGGAGTACCCGTCAAAGAGG |
| Cloning | MpCCD8-F | GAACAGCGTGAARCAGGAAC |
| | MpCCD8-R | GGRTTGGCTGTGTGTTCACAACA |
| | MpaKAI2A_cDNA_F | ATCACCATCACCATATGTCTATCCTTGAGGCTCAC |
| | MpaKAI2A_cDNA_R | GGTGGTGGTGCTCGAGCTAAATAGAACCCATAATA |
| | MpaKAI2B_cDNA_F | ATCACCATCACCATATGTCTAACCTCGAAGTTCAT |
| | MpaKAI2B_cDNA_R | GGTGGTGGTGCTCGAGTCAGCTCATTGCCACGTGC |
| | MpaMAX1_cDNA_F | GAGTTGCAGTTCTGAACGTG |
| | MpaMAX1_cDNA_R | CTGTAATCCTGTTCGTTCGTC |
| | MpaMAX1_cloning_F | CTAAATTACCGGATCATGATAACGAGCATGGAAGGATG |
| | MpaMAX1_cloning_R | GCGAATTCGAGCTCGCTACAAGCGTGTCCATCTGG |

**Supplementary Table 1. Species used in the Phylogenetic analysis.**

| abbreviation | Species name | Group | Order | Family | Source | AM symbiosis |
|---|---|---|---|---|---|---|
| Abilas | Abies lasiocarpa | Gymnosperms | Pinales | Pinaceae | 1KP | 0 |
| Acmpan | Acmopyle pancheri | Gymnosperms | Pinales | Podocarpaceae | 1KP | ND |
| Adiale | Adiantum aleuticum | Leptosporangiate | Polypodiales | Pteridaceae | 1KP | 1 |
| Adirad | Adiantum raddianum | Leptosporangiate | Polypodiales | Pteridaceae | 1KP | 1 |
| Agamac | Agathis macrophylla | Gymnosperms | Pinales | Araucariaceae | 1KP | 1 |
| Agarob | Agathis robusta | Gymnosperms | Pinales | Araucariaceae | 1KP | 1 |
| Alnglu | Alnus glutinosa | Angiosperms | Fagales | Betulaceae | 10.1126/science.aat1743 | 1 |
| Ambtri | Amborella trichopoda | Angiosperms | Amborellales | Amborellaceae | 10.1126/science.1241089 | 1 |
| Amearg | Amentotaxus argotaenia | Gymnosperms | Pinales | Taxaceae | 1KP | 1 |
| Anacom | Ananas comosus | Angiosperms | Bromeliales | Bromeliaceae | 10.1038/ng.3435 | 1 |
| Andrup | Andreaea rupestris | Mosses | Andreaeales | Andreaeaceae | 1KP | 0 |
| Anetom | Anemia tomentosa | Leptosporangiate | Schizaeales | Anemiaceae | 1KP | 1 |
| Angeve | Angiopteris evecta | Eusporangiate | Marattiales | Marattiaceae | 1KP | 1 |
| Anoatt | Anomodon attenuatus | Mosses | Hypnales | Anomodontaceae | 1KP | 0 |
| Antang | Anthoceros angustus | Hornworts | Anthocerotales | Anthocerotaceae | 1KP | 1 |
| Aposhe | Apostasia shenzhenica | Angiosperms | Asparagales | Orchidaceae | 10.1038/nature23897 | 0 |
| Arahal | Arabidopsis halleri | Angiosperms | Brassicales | Brassicaceae | Unpublished - Phytozome | 0 |
| Aralyr | Arabidopsis lyrata | Angiosperms | Brassicales | Brassicaceae | 10.1038/ng.807 | 0 |
| Aratha | Arabidopsis thaliana | Angiosperms | Brassicales | Brassicaceae | 10.1093/nar/gkr1090 | 0 |
| Aradur | Arachis duranensis | Angiosperms | Fabales | Fabaceae | 10.1038/ng.3517 | 1 |
| Arahyp | Arachis hypogaea | Angiosperms | Fabales | Fabaceae | Unpublished - peanutbase.org | 1 |
| Araipa | Arachis ipaensis | Angiosperms | Fabales | Fabaceae | 10.1038/ng.3517 | 1 |
| Ararul | Araucaria rulei | Gymnosperms | Pinales | Araucariaceae | 1KP | 1 |
| Argniv | Argyrochosma nivea | Leptosporangiate | Polypodiales | Pteridaceae | 1KP | ND |
| Aspnid | Asplenium nidus | Leptosporangiate | Polypodiales | Aspleniaceae | 1KP | 0 |
| Asppla | Asplenium platyneuron | Leptosporangiate | Polypodiales | Aspleniaceae | 1KP | 1 |
| Athcup | Athrotaxis cupressoides | Gymnosperms | Pinales | Cupressaceae | 1KP | 1 |
| Athfil | Athyrium filix-femina | Leptosporangiate | Polypodiales | Dryopteridaceae | 1KP | 1 |
| Atrang | Atrichum angustatum | Mosses | Polytrichales | Polytrichaceae | 1KP | 0 |
| Aulhet | Aulacomnium heterostichum | Mosses | Aulacomniales | Aulacomniaceae | 1KP | 0 |
| Auschi | Austrocedrus chilensis | Gymnosperms | Pinales | Cupressaceae | 1KP | 1 |
| Ausspi | Austrotaxus spicata | Gymnosperms | Pinales | Taxaceae | 1KP | 1 |
| Azocar | Azolla cf. caroliniana | Leptosporangiate | Salviniales | Salviniaceae | 1KP | 0 |
| Azofil | Azolla filiculoides | Leptosporangiate | Salviniales | Salviniaceae | 10.1038/s41477-018-0188-8 | 0 |
| Barbar | Barbilophozia barbata | Liverworts | Jungermanniales | Anastrophyllaceae | 1KP | 0 |
| Baztri | Bazzania trilobata | Liverworts | Jungermanniales | Lepidoziaceae | 1KP | 0 |
| Begfuc | Begonia fuchsioides | Angiosperms | Cucurbitales | Begoniaceae | 10.1126/science.aat1743 | 1 |
| Betvul | Beta vulgaris ssp. vulgaris KWS2320 | Angiosperms | Caryophyllales | Amaranthaceae | 10.1038/nature12817 | 0 |
| Blasp. | Blasia sp. | Liverworts | Blasiales | Blasiaceae | 1KP | 0 |
| Blespi | Blechnum spicant | Leptosporangiate | Polypodiales | Blechnaceae | 1KP | 1 |
| Boestr | Boechera stricta | Angiosperms | Brassicales | Brassicaceae | Unpublished - Phytozome | 0 |
| Bolrep | Bolbitis repanda | Leptosporangiate | Polypodiales | Dryopteridaceae | 1KP | 0 |
| Botvir | Botrypus virginianus | Eusporangiate | Ophioglossales | Ophioglossaceae | 1KP | 1 |
| Bradis | Brachypodium distachyon | Angiosperms | Poales | Poaceae | 10.1038/nature08747 | 1 |
| Braolecap | Brassica oleraceae capitata | Angiosperms | Brassicales | Brassicaceae | 10.1038/ncomms4930 | 0 |
| Brarap | Brassica rapa FPsc | Angiosperms | Brassicales | Brassicaceae | Unpublished - Phytozome | 0 |
| Bryarg | Bryum argenteum | Mosses | Bryales | Bryaceae | 1KP | 0 |
| Buxaph | Buxbaumia aphylla | Mosses | Buxbaumiales | Buxbaumiaceae | 1KP | 0 |
| Cajcaj | Cajanus cajan | Angiosperms | Fabales | Fabaceae | 10.1038/nbt.2022. | 1 |
| Calcor | Calliergon cordifolium | Mosses | Hypnales | Calliergonaceae | 1KP | 0 |
| Calgra | Callitris gracilis | Gymnosperms | Pinales | Cupressaceae | 1KP | 1 |
| Calmac | Callitris macleayana | Gymnosperms | Pinales | Cupressaceae | 1KP | 1 |
| Caldec | Calocedrus decurrens | Gymnosperms | Pinales | Cupressaceae | 1KP | 1 |
| Calfis | Calypogeia fissa | Liverworts | Jungermanniales | Calypogeiaceae | 1KP | 0 |
| Capgra | Capsella grandiflora | Angiosperms | Brassicales | Brassicaceae | 10.1038/ng.2669 | 0 |
| Caprub | Capsella rubella | Angiosperms | Brassicales | Brassicaceae | 10.1038/ng.2669 | 0 |
| Capann | Capsicum annuum cvCM334 | Angiosperms | Solanales | Solanaceae | 10.1038/ng.2877 | 1 |
| Carpap | Carica papaya | Angiosperms | Brassicales | Caricaceae | 10.1038/nature06856 | 1 |
| Casmol | Castanea mollissima | Angiosperms | Fagales | Fagaceae | unpublished | 1 |
| Casaus | Castanospermum australe | Angiosperms | Fabales | Fabaceae | 10.1126/science.aat1743 | 1 |
| Casgla | Casuarina glauca | Angiosperms | Fagales | Casuarinaceae | 10.1126/science.aat1743 | 1 |
| Catagr | Cathaya argyrophylla | Gymnosperms | Pinales | Pinaceae | 1KP | 0 |
| Cedlib | Cedrus libani | Gymnosperms | Pinales | Pinaceae | 1KP | 0 |
| Cephar | Cephalotaxus harringtonia | Gymnosperms | Pinales | Cephalotaxaceae | 1KP | 1 |
| Cepfol | Cephalotus follicularis | Angiosperms | Oxalidales | Cephalotaceae | 10.1038/s41559-016-0059 | 0 |
| Cerpur | Ceratodon purpureus | Mosses | Dicranales | Dicranaceae | 1KP | 0 |
| Cercan | Cercis canadensis | Angiosperms | Fabales | Caesalpiniaceae | 10.1126/science.aat1743 | 1 |
| Chafas | Chamaecrista fasciculata | Angiosperms | Fabales | Fabaceae | 10.1126/science.aat1743 | 1 |
| Chalaw | Chamaecyparis lawsoniana | Gymnosperms | Pinales | Cupressaceae | 1KP | 1 |

| | | | | | | |
|---|---|---|---|---|---|---|
| Chequi | Chenopodium quinoa | Angiosperms | Caryophyllales | Amaranthaceae | 10.1038/nature21370 | 0 |
| Cicari | Cicer arietinum ICC4958 | Angiosperms | Fabales | Fabaceae | 10.1038/srep12806 | 1 |
| Citlan | Citrullus lanatus subsp. vulgaris 97103 | Angiosperms | Cucurbitales | Cucurbitaceae | 10.1038/ng.2470 | 1 |
| Citcle | Citrus clementina | Angiosperms | Sapindales | Rutaceae | 10.1038/nbt.2906 | 1 |
| Citsin | Citrus sinensis | Angiosperms | Sapindales | Rutaceae | 10.1038/nbt.2906 | 1 |
| Anocla | Claopodium rostratum | Mosses | Hypnales | Hypnaceae | 1KP | 0 |
| Cliden | Climacium dendroides | Mosses | Hypnales | Climaciaceae | 1KP | 0 |
| Concon | Conocephalum conicum | Liverworts | Marchantiales | Conocephalaceae | 1KP | 1 |
| Creven | Crepidomanes venosum | Leptosporangiate | Hymenophyllales | Hymenophyllaceae | 1KP | 0 |
| Cryacr | Cryptogramma acrostichoides | Leptosporangiate | Polypodiales | Pteridaceae | 1KP | 0 |
| Cryjap | Cryptomeria japonica | Gymnosperms | Pinales | Cupressaceae | 1KP | 1 |
| Cucmel | Cucumis melo | Angiosperms | Cucurbitales | Cucurbitaceae | 10.1073/pnas.1205415109 | 1 |
| Cucsat | Cucumis sativus PI183967 | Angiosperms | Cucurbitales | Cucurbitaceae | 10.1038/ng.2801 | 1 |
| Cucmax | Cucurbita maxima | Angiosperms | Cucurbitales | Cucurbitaceae | 10.1016/j.molp.2017.09.003 | 1 |
| Cucmos | Cucurbita moschata | Angiosperms | Cucurbitales | Cucurbitaceae | 10.1016/j.molp.2017.09.004 | 1 |
| Cucpep | Cucurbita pepo | Angiosperms | Cucurbitales | Cucurbitaceae | Unpublished - cucurbitgenomics.o | 1 |
| Culmac | Culcita macrocarpa | Leptosporangiate | Cyatheales | Culcitaceae | 1KP | 1 |
| Cunlan | Cunninghamia lanceolata | Gymnosperms | Pinales | Cupressaceae | 1KP | 1 |
| Cupdup | Cupressus dupreziana | Gymnosperms | Pinales | Cupressaceae | 1KP | 1 |
| Cyaspi | Cyathea (Alsophila) spinulosa | Leptosporangiate | Cyatheales | Cyatheaceae | 1KP | 1 |
| Cycmic | Cycas micholitzii | Gymnosperms | Cycadales | Cycadaceae | Unpublished - gymno-plaza | 1 |
| Cycmic | Cycas micholitzii | Gymnosperms | Cycadales | Cycadaceae | 1KP | 1 |
| Cysfra | Cystopteris fragilis | Leptosporangiate | Polypodiales | Cystopteridaceae | 1KP | 1 |
| Cyspro | Cystopteris protrusa | Leptosporangiate | Polypodiales | Cystopteridaceae | 1KP | 1 |
| Cysree | Cystopteris reevesiana | Leptosporangiate | Polypodiales | Cystopteridaceae | 1KP | 1 |
| Cysuta | Cystopteris utahensis | Leptosporangiate | Polypodiales | Cystopteridaceae | 1KP | 1 |
| Daccom | Dacrycarpus compactus | Gymnosperms | Pinales | Podocarpaceae | 1KP | 1 |
| Dacbal | Dacrydium balansae | Gymnosperms | Pinales | Podocarpaceae | 1KP | 1 |
| Datglo | Datisca glomerata | Angiosperms | Cucurbitales | Dasticaceae | 10.1126/science.aat1743 | 1 |
| Daucar | Daucus carota | Angiosperms | Apiales | Apiaceae | 10.1038/ng.3565 | 1 |
| Davfej | Davallia fejeensis | Leptosporangiate | Polypodiales | Davalliaceae | 1KP | ND |
| Dencat | Dendrobium catenatum | Angiosperms | Asparagales | Orchidaceae | 10.1038/nature23897 | 0 |
| Denobs | Dendrolycopodium obscurum | Lycophytes | Lycopodiales | Lycopodiaceae | 1KP | 0 |
| Dendav | Dennstaedtia davallioides | Leptosporangiate | Polypodiales | Dennstaedtiaceae | 1KP | 1 |
| Deplob | Deparia lobato-crenata | Leptosporangiate | Polypodiales | Athyriaceae | 1KP | ND |
| Diacar | Dianthus caryophyllus | Angiosperms | Caryophyllales | Caryophyllaceae | 10.1093/dnares/dst053 | 0 |
| Dicsco | Dicranum scoparium | Mosses | Dicranales | Dicranaceae | 1KP | 0 |
| Didtru | Didymochlaena truncatula | Leptosporangiate | Polypodiales | Hypodematiaceae | 1KP | ND |
| Dioedu | Dioon edule | Gymnosperms | Cycadales | Zamiaceae | 1KP | 1 |
| Dipdig | Diphasiastrum digitatum | Lycophytes | Lycopodiales | Lycopodiaceae | 1KP | 1 |
| Dipfol | Diphyscium foliosum | Mosses | Diphysciales | Diphysciaceae | 1KP | 0 |
| Dipwic | Diplazium wichurae | Leptosporangiate | Polypodiales | Athyriaceae | 1KP | 1 |
| Dipcon | Dipteris conjugata | Leptosporangiate | Gleicheniales | Dipteridaceae | 1KP | ND |
| Distri | Discaria trinervis | Angiosperms | Rosales | Rhamnaceae | 10.1126/science.aat1743 | 1 |
| Disarc | Diselma archeri | Gymnosperms | Pinales | Cupressaceae | 1KP | 1 |
| Drydru | Dryas drummondii | Angiosperms | Rosales | Rosaceae | 10.1126/science.aat1743 | 1 |
| Encstr | Encalypta streptocarpa | Mosses | Funariales | Encalyptaceae | 1KP | 0 |
| Encbar | Encephalartos barteri | Gymnosperms | Cycadales | Zamiaceae | 1KP | 1 |
| Ephsin | Ephedra sinica | Gymnosperms | Ephedrales | Ephedraceae | 1KP | 1 |
| Equdif | Equisetum diffusum | Eusporangiate | Equisetales | Equisetaceae | 1KP | 1 |
| Equhye | Equisetum hyemale | Eusporangiate | Equisetales | Equisetaceae | 1KP | 1 |
| Eutsal | Eutrema salsugineum | Angiosperms | Brassicales | Brassicaceae | 10.3389/fpls.2013.00046 | 0 |
| Faltax | Falcatifolium taxoides | Gymnosperms | Pinales | Podocarpaceae | 1KP | 1 |
| Fokhod | Fokienia hodginsii | Gymnosperms | Pinales | Cupressaceae | 1KP | 1 |
| Fonant | Fontinalis antipyretica | Mosses | Isobryales | Fontinalaceae | 1KP | 0 |
| Fraves | Fragaria vesca | Angiosperms | Rosales | Rosaceae | 10.1093/gigascience/gix124 | 1 |
| Fraana | Fragaria x ananassa | Angiosperms | Rosales | Rosaceae | Unpublished - rosaceae.org | 1 |
| Fraexc | Fraxinus excelsior | Angiosperms | Lamiales | Oleaceae | doi:10.1038/nature20786 | 1 |
| Frusp. | Frullania sp. | Liverworts | Porellales | Frullaniaceae | 1KP | ND |
| Fruspp | Frullania spp. | Liverworts | Porellales | Frullaniaceae | 1KP | ND |
| Gagari | Gaga arizonica | Leptosporangiate | Polypodiales | Pteridaceae | 1KP | ND |
| Ginbil | Ginkgo biloba | Gymnosperms | Ginkgoales | Ginkgoaceae | 10.1186/s13742-016-0154-1 | 1 |
| Ginbil | Ginkgo biloba | Gymnosperms | Ginkgoales | Ginkgoaceae | 1KP | 1 |
| Glymax | Glycine max | Angiosperms | Fabales | Fabaceae | 10.1038/nature08670 | 1 |
| Glypen | Glyptostrobus pensilis | Gymnosperms | Pinales | Cupressaceae | 1KP | 1 |
| Gnemon | Gnetum montanum | Gymnosperms | Gnetales | Gnetaceae | 1KP | 0 |
| Gnemon | Gnetum montanum | Gymnosperms | Gnetales | Gnetaceae | 10.5061/dryad.0vm37 | 0 |
| Gosrai | Gossypium raimondii | Angiosperms | Malvales | Malvaceae | 10.1038/nature11798 | 1 |
| Gymdry | Gymnocarpium dryopteris | Leptosporangiate | Polypodiales | Cystopteridaceae | 1KP | 1 |
| Halbid | Halocarpus bidwillii | Gymnosperms | Pinales | Podocarpaceae | 1KP | 1 |
| Hedcil | Hedwigia ciliata | Mosses | Hedwigiales | Hedwigiaceae | 1KP | 0 |
| Helann | Helianthus annuus | Angiosperms | Asterales | Asteraceae | 10.1038/nature22380 | 1 |

| | | | | | | |
|---|---|---|---|---|---|---|
| Hompyc | Homalosorus pycnocarpos | Leptosporangiate | Polypodiales | Aspleniaceae | 1KP | ND |
| Horvul | Hordeum vulgare | Angiosperms | Poales | Poaceae | 10.1038/nature22043 / 10.1038/s | 1 |
| Humlup | Humulus lupulus | Angiosperms | Rosales | Cannabaceae | 10.1093/pcp/pcu169 | 1 |
| Hupluc | Huperzia lucidula | Lycophytes | Lycopodiales | Lycopodiaceae | 1KP | 1 |
| Hupmyr | Huperzia myrsinites | Lycophytes | Lycopodiales | Lycopodiaceae | 1KP | 1 |
| Hupsel | Huperzia selago | Lycophytes | Lycopodiales | Lycopodiaceae | 1KP | 1 |
| Hupsqu | Huperzia squarrosa | Lycophytes | Lycopodiales | Lycopodiaceae | 1KP | 1 |
| Hymbiv | Hymenophyllum bivalve | Leptosporangiate | Hymenophyllales | Hymenophyllaceae | 1KP | 0 |
| Hymcup | Hymenophyllum cupressiforme | Leptosporangiate | Hymenophyllales | Hymenophyllaceae | 1KP | 0 |
| Isoesp. | Isoetes sp. | Lycophytes | Isoëtales | Isoëtaceae | 1KP | ND |
| Isoteg | Isoetes tegetiformans | Lycophytes | Isoëtales | Isoëtaceae | 1KP | ND |
| Jugreg | Juglans regia | Angiosperms | Fagales | Juglandaceae | 10.1111/tpj.13207 | 1 |
| Junsco | Juniperus scopulorum | Gymnosperms | Pinales | Cupressaceae | 1KP | 1 |
| Keteve | Keteleeria evelyniana | Gymnosperms | Pinales | Pinaceae | 1KP | 0 |
| Lagfra | Lagarostrobos franklinii | Gymnosperms | Pinales | Podocarpaceae | 1KP | 1 |
| Lagsic | Lagenaria siceraria | Angiosperms | Cucurbitales | Cucurbitaceae | 10.1111/tpj.13722 | 1 |
| Larspe | Larix speciosa | Gymnosperms | Pinales | Pinaceae | 1KP | 0 |
| Leidus | Leiosporoceros dussii | Hornworts | Leiosporocerotales | Leiosporocerotacea | 1KP | 0 |
| Lepsp. | Lepidothamnus sp. | Gymnosperms | Pinales | Podocarpaceae | 1KP | 1 |
| Leualb | Leucobryum albidum | Mosses | Dicranales | Leucobryaceae | 1KP | 0 |
| Leugla | Leucobryum glaucum | Mosses | Dicranales | Leucobryaceae | 1KP | 0 |
| Leubra | Leucodon brachypus | Mosses | Hypnales | Leucodontaceae | 1KP | 0 |
| Leujul | Leucodon julaceus | Mosses | Hypnales | Leucodontaceae | 1KP | 0 |
| Leuimm | Leucostegia immersa | Leptosporangiate | Polypodiales | Hypodematiaceae | 1KP | ND |
| Linlin | Lindsaea linearis | Leptosporangiate | Polypodiales | Lindsaeaceae | 1KP | 1 |
| Linmic | Lindsaea microphylla | Leptosporangiate | Polypodiales | Lindsaeaceae | 1KP | 1 |
| Loebre | Loeskeobryum brevirostre | Mosses | Hypnales | Hylocomiaceae | 1KP | 0 |
| Lotjap | Lotus japonicus | Angiosperms | Fabales | Fabaceae | 10.1093/dnares/dsn008 | 1 |
| Luncru | Lunularia cruciata | Liverworts | Marchantiales | Lunulariaceae | 1KP | 1 |
| Lupang | Lupinus angustifolius | Angiosperms | Fabales | Fabaceae | 10.1111/pbi.12615 | 0 |
| Lycapp | Lycopodiella appressa | Lycophytes | Lycopodiales | Lycopodiaceae | 1KP | 1 |
| Lycann | Lycopodium annotinum | Lycophytes | Lycopodiales | Lycopodiaceae | 1KP | 1 |
| Lycdeu | Lycopodium deuterodensum | Lycophytes | Lycopodiales | Lycopodiaceae | 1KP | 1 |
| Lygjap | Lygodium japonicum | Leptosporangiate | Schizaeales | Lygodiaceae | 1KP | 1 |
| Maldom | Malus domestica | Angiosperms | Rosales | Rosaceae | Unpublished - rosaceae.org | 1 |
| Manesc | Manihot esculenta | Angiosperms | Malpighiales | Euphorbiaceae | 10.1038/nbt.3535 | 1 |
| Mancol | Manoao colensoi | Gymnosperms | Pinales | Podocarpaceae | 1KP | 1 |
| Marsp. | Marattia sp. | Eusporangiate | Marattiales | Marattiaceae | 1KP | 1 |
| Marema | Marchantia emarginata | Liverworts | Marchantiales | Marchantiaceae | 1KP | 1 |
| Marpal | Marchantia paleacea | Liverworts | Marchantiales | Marchantiaceae | This study | 1 |
| Marpal | Marchantia paleacea | Liverworts | Marchantiales | Marchantiaceae | 1KP | 1 |
| Marpol | Marchantia polymorpha | Liverworts | Marchantiales | Marchantiaceae | 10.1016/j.cell.2017.09.030 | 0 |
| Marpol | Marchantia polymorpha | Liverworts | Marchantiales | Marchantiaceae | 1KP | 0 |
| Medtru | Medicago truncatula | Angiosperms | Fabales | Fabaceae | 10.1038/s41477-018-0286-7 | 1 |
| Megfla | Megaceros flagellaris | Hornworts | Dendrocerotales | Dendrocerotaceae | 1KP | 0 |
| Metgly | Metasequoia glyptostroboides | Gymnosperms | Pinales | Cupressaceae | 1KP | 1 |
| Metcra | Metzgeria crassipilis | Liverworts | Metzgeriales | Metzgeriaceae | 1KP | 0 |
| Micdec | Microbiota decussata | Gymnosperms | Pinales | Cupressaceae | 1KP | 1 |
| Mictet | Microcachrys tetragona | Gymnosperms | Pinales | Podocarpaceae | 1KP | 1 |
| Mimpud | Mimosa pudica | Angiosperms | Fabales | Fabaceae | 10.1126/science.aat1743 | 1 |
| Mixspe | mixed species | Liverworts | NA | NA | 1KP | ND |
| Momcha | Momordica charantia | Angiosperms | Cucurbitales | Cucurbitaceae | Unpublished - NCBI | 1 |
| Mongot | Monoclea gottschei | Liverworts | Marchantiales | Monocleaceae | 1KP | 1 |
| Mornot | Morus notabilis | Angiosperms | Rosales | Moraceae | 10.1038/ncomms3445 | 1 |
| Musacu | Musa acuminata | Angiosperms | Zingiberales | Zingiberaceae | 10.1093/database/bat035 | 1 |
| Myrruf | Myriopteris rufa | Leptosporangiate | Polypodiales | Pteridaceae | 1KP | 1 |
| Nagnag | Nageia nagi | Gymnosperms | Pinales | Podocarpaceae | 1KP | 1 |
| Necdou | Neckera douglasii | Mosses | Hypnales | Neckeraceae | 1KP | 0 |
| Nelnuc | Nelumbo nucifera | Angiosperms | Proteales | Nelumbonaceae | 10.1186/gb-2013-14-5-r41 | 0 |
| Neopan | Neocallitropsis pancheri | Gymnosperms | Pinales | Cupressaceae | 1KP | 1 |
| Nepexa | Nephrolepis exaltata | Leptosporangiate | Polypodiales | Nephrolepidaceae | 1KP | 1 |
| Nicben | Nicotiana benthamiana | Angiosperms | Solanales | Solanaceae | 10.1094/MPMI-06-12-0148-TA | 1 |
| Nissch | Nissolia schottii | Angiosperms | Fabales | Fabaceae | 10.1126/science.aat1743 | 1 |
| Notaen | Nothoceros aenigmaticus | Hornworts | Dendrocerotales | Dendrocerotaceae | 1KP | ND |
| Notvin | Nothoceros vincentianus | Hornworts | Dendrocerotales | Dendrocerotaceae | 1KP | 1 |
| Notmon | Notholaena montieliae | Leptosporangiate | Polypodiales | Pteridaceae | 1KP | ND |
| Notlon | Nothotsuga longibracteata | Gymnosperms | Pinales | Pinaceae | 1KP | 0 |
| Odopro | Odontoschisma prostratum | Liverworts | Jungermanniales | Cephaloziaceae | 1KP | 0 |
| Onosen | Onoclea sensibilis | Leptosporangiate | Polypodiales | Onocleaceae | 1KP | 1 |
| Ophpet | Ophioglossum petiolatum | Eusporangiate | Ophioglossales | Ophioglossaceae | 1KP | 1 |
| Ophvul | Ophioglossum vulgatum | Eusporangiate | Ophioglossales | Ophioglossaceae | 1KP | 1 |
| Ortlye | Orthotrichum lyellii | Mosses | Orthotrichales | Orthotrichaceae | 1KP | 0 |

| | | | | | | |
|---|---|---|---|---|---|---|
| Orysat | Oryza sativa | Angiosperms | Poales | Poaceae | 10.1093/nar/gkl976 | 1 |
| Osmjav | Osmunda javanica | Leptosporangiate | Osmundales | Osmundaceae | 1KP | 1 |
| Osmreg | Osmunda regalis | Leptosporangiate | Osmundales | Osmundaceae | 1KP | 1 |
| Osmsp. | Osmunda sp. | Leptosporangiate | Osmundales | Osmundaceae | 1KP | 1 |
| Osmcin | Osmundastrum cinnamomeum | Leptosporangiate | Osmundales | Osmundaceae | 1KP | 1 |
| Pallye | Pallavicinia lyellii | Liverworts | Pallaviciniales | Pallaviciniaceae | 1KP | 1 |
| Pappap | Papuacedrus papuana | Gymnosperms | Pinales | Cupressaceae | 1KP | 1 |
| Parcor | Parahemionitis cordata | Leptosporangiate | Polypodiales | Pteridaceae | 1KP | 1 |
| Parhal | Paraphymatoceros hallii | Hornworts | Notothyladales | Notothyladaceae | 1KP | 0 |
| Parust | Parasitaxus usta | Gymnosperms | Pinales | Podocarpaceae | 1KP | ND |
| Parand | Parasponia andersonii | Angiosperms | Rosales | Cannabaceae | 10.1073/pnas.1721395115 | 1 |
| Pelepi | Pellia cf. epiphylla | Liverworts | Pelliales | Pelliaceae | 1KP | 1 |
| Pelnee | Pellia neesiana | Liverworts | Pelliales | Pelliaceae | 1KP | 1 |
| Petaxi | Petunia axillaris | Angiosperms | Solanales | Solanaceae | 10.1038/nplants.2016.74 | 1 |
| Phacar | Phaeoceros carolinianus | Hornworts | Notothyladales | Notothyladaceae | 1KP | 1 |
| Phaequ | Phalaenopsis equestris | Angiosperms | Asparagales | Orchidaceae | 10.1038/nature23897 | 0 |
| Phavul | Phaseolus vulgaris | Angiosperms | Fabales | Fabaceae | 10.1038/ng.3008 | 1 |
| Phifon | Philonotis fontana | Mosses | Bartramiales | Bartramiaceae | 1KP | 0 |
| Phlpse | Phlebodium pseudoaureum | Leptosporangiate | Polypodiales | Polypodiaceae | 1KP | ND |
| Phyhyp | Phyllocladus hypophyllus | Gymnosperms | Pinales | Podocarpaceae | 1KP | 1 |
| Phydru | Phylloglossum drummondii | Lycophytes | Lycopodiales | Lycopodiaceae | 1KP | 1 |
| Phygro | Phymatosorus grossus | Leptosporangiate | Polypodiales | Polypodiaceae | 1KP | 0 |
| Phypat | Physcomitrella patens | Mosses | Funariales | Funariaceae | 10.1111/tpj.13801 | 0 |
| Physsp. | Physcomitrium sp. | Mosses | Funariales | Physcomitrium | 1KP | 0 |
| Picabi | Picea abies | Gymnosperms | Pinales | Pinaceae | 10.1038/nature12211 | 0 |
| Piceng | Picea engelmannii | Gymnosperms | Pinales | Pinaceae | 1KP | 0 |
| Picgla | Picea glauca | Gymnosperms | Pinales | Pinaceae | 10.1093/bioinformatics/btt178 | 0 |
| Picsit | Picea sitchensis | Gymnosperms | Pinales | Pinaceae | Unpublished - gymno-plaza | 0 |
| Piluvi | Pilgerodendron uviferum | Gymnosperms | Pinales | Cupressaceae | 1KP | 1 |
| Pilglo | Pilularia globulifera | Leptosporangiate | Salviniales | Marsileaceae | 1KP | 0 |
| Pinjef | Pinus jeffreyi | Gymnosperms | Pinales | Pinaceae | 1KP | 0 |
| Pinpar | Pinus parviflora | Gymnosperms | Pinales | Pinaceae | 1KP | 0 |
| Pinpin | Pinus pinaster | Gymnosperms | Pinales | Pinaceae | Unpublished - gymno-plaza | 0 |
| Pinpon | Pinus ponderosa | Gymnosperms | Pinales | Pinaceae | 1KP | 0 |
| Pinrad | Pinus radiata | Gymnosperms | Pinales | Pinaceae | 1KP | 0 |
| Pinsyl | Pinus sylvestris | Gymnosperms | Pinales | Pinaceae | Unpublished - gymno-plaza | 0 |
| Pintae | Pinus taeda | Gymnosperms | Pinales | Pinaceae | 10.1534/genetics.113.159715 | 0 |
| Pittri | Pityrogramma trifoliata | Leptosporangiate | Polypodiales | Pteridaceae | 1KP | 1 |
| Plajap | Plagiogyria japonica | Leptosporangiate | Cyatheales | Plagiogyriaceae | 1KP | 1 |
| Plains | Plagiomnium insigne | Mosses | Bryales | Mniaceae | 1KP | 0 |
| Plaori | Platycladus orientalis | Gymnosperms | Pinales | Cupressaceae | 1KP | 1 |
| Plepol | Pleopeltis polypodioides | Leptosporangiate | Polypodiales | Polypodiaceae | 1KP | 0 |
| Podcor | Podocarpus coriaceus | Gymnosperms | Pinales | Podocarpaceae | 1KP | 1 |
| Podrub | Podocarpus rubens | Gymnosperms | Pinales | Podocarpaceae | 1KP | 1 |
| Polamo | Polypodium amorphum | Leptosporangiate | Polypodiales | Polypodiaceae | 1KP | ND |
| Polgly | Polypodium glycyrrhiza | Leptosporangiate | Polypodiales | Polypodiaceae | 1KP | 1 |
| Polhes | Polypodium hesperium | Leptosporangiate | Polypodiales | Polypodiaceae | 1KP | ND |
| Polacr | Polystichum acrostichoides | Leptosporangiate | Polypodiales | Dryopteridaceae | 1KP | 1 |
| Polcom | Polytrichum commune | Mosses | Polytrichales | Polytrichaceae | 1KP | 0 |
| Poptri | Populus trichocarpa | Angiosperms | Malpighiales | Salicaceae | 10.1126/science.1128691 | 1 |
| Pornav | Porella navicularis | Liverworts | Porellales | Porellaceae | 1KP | ND |
| Porpin | Porella pinnata | Liverworts | Porellales | Porellaceae | 1KP | ND |
| Potmic | Potentilla micrantha | Angiosperms | Rosales | Rosaceae | 10.1093/gigascience/giy010 | 1 |
| Pruand | Prumnopitys andina | Gymnosperms | Pinales | Podocarpaceae | 1KP | 0 |
| Pruavi | Prunus avium | Angiosperms | Rosales | Rosaceae | 10.1093/dnares/dsx020 | 1 |
| Prudul | Prunus dulcis | Angiosperms | Rosales | Rosaceae | Unpublished - rosaceae.org | 1 |
| Prumum | Prunus mume | Angiosperms | Rosales | Rosaceae | Unpublished - NCBI | 1 |
| Pruper | Prunus persica | Angiosperms | Rosales | Rosaceae | 10.1038/ng.2586 | 1 |
| Pseama | Pseudolarix amabilis | Gymnosperms | Pinales | Pinaceae | 1KP | 0 |
| Psecar | Pseudolycopodiella caroliniana | Lycophytes | Lycopodiales | Lycopodiaceae | 1KP | ND |
| Pseele | Pseudotaxiphyllum elegans | Mosses | Hypnales | Hypnaceae | 1KP | 0 |
| Psechi | Pseudotaxus chienii | Gymnosperms | Pinales | Taxaceae | 1KP | 1 |
| Psemen | Pseudotsuga menziesii | Gymnosperms | Pinales | Pinaceae | 10.1534/g3.117.300078 | 0 |
| Psewil | Pseudotsuga wilsoniana | Gymnosperms | Pinales | Pinaceae | 1KP | 0 |
| Psinud | Psilotum nudum | Eusporangiate | Psilotales | Psilotaceae | 1KP | 1 |
| Pteens | Pteris ensiformis | Leptosporangiate | Polypodiales | Pteridaceae | 1KP | 1 |
| Ptevit | Pteris vittata | Leptosporangiate | Polypodiales | Pteridaceae | 1KP | 1 |
| Ptipul | Ptilidium pulcherrimum | Liverworts | Ptilidiales | Ptilidiaceae | 1KP | ND |
| Pyrcom | Pyrus communis | Angiosperms | Rosales | Rosaceae | 10.1371/journal.pone.0092644 | 1 |
| Pyrbre | Pyrus x bretschneideri | Angiosperms | Rosales | Rosaceae | Unpublished - NCBI | 1 |
| Querob | Quercus robur | Angiosperms | Fagales | Fagaceae | 10.1111/1755-0998.12425 | 1 |
| Racelo | Racomitrium elongatum | Mosses | Grimmiales | Grimmiaceae | 1KP | 0 |

| | | | | | | |
|---|---|---|---|---|---|---|
| Racvar | Racomitrium varium | Mosses | Grimmiales | Grimmiaceae | 1KP | 0 |
| Radlin | Radula lindenbergiana | Liverworts | Porellales | Radulaceae | 1KP | ND |
| Retmin | Retrophyllum minus | Gymnosperms | Pinales | Podocarpaceae | 1KP | 1 |
| Rhyser | Rhynchostegium serrulatum | Mosses | Hypnales | Brachytheciaceae | 1KP | 0 |
| Ricnat | Ricciocarpos natans | Liverworts | Marchantiales | Ricciaceae | 1KP | 0 |
| Riccom | Ricinus communis | Angiosperms | Malpighiales | Euphorbiaceae | 10.1038/nbt.1674 | 1 |
| Roschi | Rosa chinensis | Angiosperms | Rosales | Rosaceae | 10.1038/s41588-018-0110-3 | 1 |
| Roscap | Rosulabryum cf. capillare | Mosses | Bryales | Bryaceae | 1KP | 0 |
| Rubocc | Rubus occidentalis | Angiosperms | Rosales | Rosaceae | 10.1111/tpj.13215 | 1 |
| Salcuc | Salvinia cucullata | Leptosporangiate | Salviniales | Salviniaceae | 10.1038/s41477-018-0188-9 | 0 |
| Saxcon | Saxegothaea conspicua | Gymnosperms | Pinales | Podocarpaceae | 1KP | 1 |
| Scanem | Scapania nemorosa | Liverworts | Jungermanniales | Scapaniaceae | 1KP | 0 |
| Scedis | Sceptridium dissectum | Eusporangiate | Ophioglossales | Ophioglossaceae | 1KP | 1 |
| Schisp. | Schistochila sp. | Liverworts | Jungermanniales | Schistochilaceae | 1KP | 0 |
| Sciver | Sciadopitys verticillata | Gymnosperms | Pinales | Sciadopityaceae | 1KP | 1 |
| Scoaqu | Scouleria aquatica | Mosses | Scouleriales | Scouleriaceae | 1KP | 0 |
| Selaca | Selaginella acanthonota | Lycophytes | Selaginellales | Selaginellaceae | 1KP | 1 |
| Selapo | Selaginella apoda | Lycophytes | Selaginellales | Selaginellaceae | 1KP | 1 |
| Selkra | Selaginella kraussiana | Lycophytes | Selaginellales | Selaginellaceae | 1KP | 1 |
| Sellep | Selaginella lepidophylla | Lycophytes | Selaginellales | Selaginellaceae | 1KP | 1 |
| Selmoe | Selaginella moellendorffii | Lycophytes | Selaginellales | Selaginellaceae | 10.1126/science.1203810 | 1 |
| Selsel | Selaginella selaginoides | Lycophytes | Selaginellales | Selaginellaceae | 1KP | 1 |
| Selsta | Selaginella stauntoniana | Lycophytes | Selaginellales | Selaginellaceae | 1KP | 1 |
| Selwal | Selaginella wallacei | Lycophytes | Selaginellales | Selaginellaceae | 1KP | 1 |
| Selwil | Selaginella willdenowii | Lycophytes | Selaginellales | Selaginellaceae | 1KP | 1 |
| Seqsem | Sequoia sempervirens | Gymnosperms | Pinales | Cupressaceae | 1KP | 1 |
| Seqgig | Sequoiadendron giganteum Glaucum | Gymnosperms | Pinales | Cupressaceae | 1KP | 1 |
| Setita | Setaria italica | Angiosperms | Poales | Poaceae | 10.1038/nbt.2196 | 1 |
| Sollyc | Solanum lycopersicum | Angiosperms | Solanales | Solanaceae | 10.1038/nature11119 | 1 |
| Solpen | Solanum pennellii | Angiosperms | Solanales | Solanaceae | 10.1038/ng.3046 | 1 |
| Sorbic | Sorghum bicolor | Angiosperms | Poales | Poaceae | 10.1111/tpj.13781 | 1 |
| Sphtex | Sphaerocarpos texanus | Liverworts | Sphaerocarpales | Sphaerocarpaceae | 1KP | 0 |
| Sphfal | Sphagnum fallax | Mosses | Sphagnales | Sphagnaceae | Unpublished - Phytozome | 0 |
| Sphles | Sphagnum lescurii | Mosses | Sphagnales | Sphagnaceae | 1KP | 0 |
| Sphpal | Sphagnum palustre | Mosses | Sphagnales | Sphagnaceae | 1KP | 0 |
| Sphrec | Sphagnum recurvum | Mosses | Sphagnales | Sphagnaceae | 1KP | 0 |
| Spiole | Spinacia oleracea | Angiosperms | Caryophyllales | Amaranthaceae | Unpublished - bvseq.molgen.mpg. | 0 |
| Spipol | Spirodela polyrhiza | Angiosperms | Alismatales | Araceae | 10.1038/ncomms4311 | 0 |
| Staeri | Stangeria eriopus | Gymnosperms | Cycadales | Zamiaceae | 1KP | 1 |
| Stesub | Stereodon subimponens | Mosses | Hypnales | Hypnaceae | 1KP | 0 |
| Stilob | Sticherus lobatus | Leptosporangiate | Gleicheniales | Gleicheniaceae | 1KP | ND |
| Sunama | Sundacarpus amarus | Gymnosperms | Pinales | Podocarpaceae | 1KP | 1 |
| Synpri | Syntrichia princeps | Mosses | Pottiales | Pottiaceae | 1KP | 0 |
| Taicry | Taiwania cryptomerioides | Gymnosperms | Pinales | Cupressaceae | 1KP | 1 |
| Taklep | Takakia lepidozioides | Mosses | Takakiales | Takakiaceae | 1KP | 1 |
| Tarhas | Tarenaya hassleriana | Angiosperms | Brassicales | Cleomaceae | 10.1105/tpc.113.113480 | 0 |
| Taxdis | Taxodium distichum | Gymnosperms | Pinales | Cupressaceae | 1KP | 1 |
| Taxbac | Taxus baccata | Gymnosperms | Pinales | Taxaceae | 1KP | 1 |
| Taxcus | Taxus cuspidata | Gymnosperms | Pinales | Taxaceae | 1KP | 1 |
| Tetsp. | Tetraclinis sp. | Gymnosperms | Pinales | Cupressaceae | 1KP | 1 |
| Tetpel | Tetraphis pellucida | Mosses | Tetraphidales | Tetraphidaceae | 1KP | 0 |
| Theacu | Thelypteris acuminata | Leptosporangiate | Polypodiales | Thelypteridaceae | 1KP | 1 |
| Thecac | Theobroma cacao | Angiosperms | Malvales | Malvaceae | 10.1186/gb-2013-14-6-r53 | 1 |
| Thudel | Thuidium delicatulum | Mosses | Hypnales | Thuidiaceae | 1KP | 0 |
| Thupli | Thuja plicata | Gymnosperms | Pinales | Cupressaceae | 1KP | 1 |
| Thudol | Thujopsis dolabrata | Gymnosperms | Pinales | Cupressaceae | 1KP | 1 |
| Thyele | Thyrsopteris elegans | Leptosporangiate | Cyatheales | Thyrsopteridaceae | 1KP | 1 |
| Timaus | Timmia austriaca | Mosses | Timmiales | Timmiaceae | 1KP | 0 |
| Tmepar | Tmesipteris parva | Eusporangiate | Psilotales | Psilotaceae | 1KP | 1 |
| Tornuc | Torreya nucifera | Gymnosperms | Pinales | Taxaceae | 1KP | 1 |
| Tortax | Torreya taxifolia | Gymnosperms | Pinales | Taxaceae | 1KP | 1 |
| Treori | Trema orientalis | Angiosperms | Rosales | Cannabaceae | 10.1073/pnas.1721395115 | 1 |
| Trelac | Treubia lacunosa | Liverworts | Treubiales | Treubiaceae | 1KP | 1 |
| Tripra | Trifolium pratense | Angiosperms | Fabales | Fabaceae | 10.1038/srep17394 | 1 |
| Trisub | Trifolium subterraneum | Angiosperms | Fabales | Fabaceae | Unpublished - NCBI | 1 |
| Tsuhet | Tsuga heterophylla | Gymnosperms | Pinales | Pinaceae | 1KP | 0 |
| Utrgib | Utricularia gibba | Angiosperms | Lamiales | Lentibulariaceae | 10.1073/pnas.1702072114 | 0 |
| Vigang | Vigna angularis | Angiosperms | Fabales | Fabaceae | 10.1038/srep080669 | 1 |
| Vigrad | Vigna radiata | Angiosperms | Fabales | Fabaceae | 10.1038/ncomms6443 | 1 |
| Vigung | Vigna unguiculata | Angiosperms | Fabales | Fabaceae | Unpublished - Phytozome | 1 |
| Vitapp | Vittaria appalachiana | Leptosporangiate | Polypodiales | Pteridaceae | 1KP | 1 |
| Vitlin | Vittaria lineata | Leptosporangiate | Polypodiales | Pteridaceae | 1KP | 1 |

| | | | | | | |
|---|---|---|---|---|---|---|
| Welmir | Welwitschia mirabilis | Gymnosperms | Gnetales | Welwitschiaceae | 1KP | 1 |
| Widced | Widdringtonia cedarbergensis | Gymnosperms | Pinales | Cupressaceae | 1KP | 1 |
| Wolnob | Wollemia nobilis | Gymnosperms | Pinales | Araucariaceae | 1KP | 1 |
| Wooilv | Woodsia ilvensis | Leptosporangiate | Polypodiales | Woodsiaceae | 1KP | 0 |
| Woosco | Woodsia scopulina | Leptosporangiate | Polypodiales | Woodsiaceae | 1KP | 0 |
| Zeamay | Zea mays PH207 | Angiosperms | Poales | Poaceae | 10.1105/tpc.16.00353 | 1 |
| Zizjuj | Ziziphus jujuba cv. Dongzao | Angiosperms | Rosales | Rhamnaceae | 10.1038/ncomms6315 | 1 |
| Zosmar | Zostera marina | Angiosperms | Alismatales | Zosteraceae | 10.1038/nature16548 | 0 |
